# Processive movement of *Staphylococcus aureus* essential septal peptidoglycan synthases is independent of FtsZ treadmilling and drives cell constriction

**DOI:** 10.1101/2023.06.29.547026

**Authors:** Simon Schäper, António D. Brito, Bruno M. Saraiva, Georgia R. Squyres, Matthew J. Holmes, Ethan C. Garner, Zach Hensel, Ricardo Henriques, Mariana G. Pinho

## Abstract

Bacterial cell division is mediated by the tubulin-homolog FtsZ, which recruits peptidoglycan (PG) synthesis enzymes to the division site. Septal PG synthases promote inward growth of the division septum, but the mechanisms governing the spatiotemporal regulation of these enzymes are poorly understood. Recent studies on various organisms have proposed different models for the relationship between the movement and activity of septum-specific PG synthases and FtsZ treadmilling. Here, we studied the movement dynamics of conserved cell division proteins relative to the rates of septum constriction and FtsZ treadmilling in the Gram-positive pathogen *Staphylococcus aureus*. The septal PG synthesis enzyme complex FtsW/PBP1 and its putative activator protein, DivIB, moved processively, around the division site, with the same velocity. Impairing FtsZ treadmilling did not affect FtsW and DivIB velocities or septum constriction rates. Contrarily, inhibition of PG synthesis slowed down or completely stopped both septum constriction and the directional movement of FtsW/PBP1 and DivIB. Our findings support a model for *S. aureus* in which a single population of processively moving FtsW/PBP1 remains associated with DivIB to drive cell constriction independently of treadmilling FtsZ filaments.

## Introduction

Bacterial cell division starts with mid-cell assembly of the divisome, a membrane-spanning multi-protein complex that is organized by the tubulin homologue FtsZ ^1^. FtsZ monomers polymerize into protofilaments that assemble along the circumference of the cytoplasmic membrane into a dynamic and patchy ring-like polymer structure, termed the Z-ring. This ring marks the future division site and acts as a scaffold to recruit other cell division proteins, including enzymes that synthesize peptidoglycan (PG), the main component of the bacterial cell wall.

FtsZ has guanosine triphosphatase (GTPase) activity, and GTP hydrolysis by FtsZ has a negative effect on the stability of protofilaments, leading to treadmilling behaviour around the division site ^2–4^. Treadmilling FtsZ filaments condense into a dense Z-ring and initiate cell constriction by guiding septal cell wall synthesis ^5, 6^. Therefore, to trigger cell constriction, PG synthases must be spatiotemporally organized and their enzymatic activities tightly regulated ^7–10^. Although *in vitro* evidence exists for FtsZ-mediated membrane constriction ^11^, PG synthesis is thought to be the main driving force for bacterial fission ^12–14^.

PG synthesis requires the activity of glycosyltransferases (GTases), which polymerise glycan strands, and transpeptidases (TPases), which cross-link glycans via peptide bridges ^15^. Unlike rod-shaped bacteria, which contain numerous Penicillin-Binding Proteins (PBPs) for peptidoglycan synthesis, *Staphylococcus aureus* encodes only four: the class A (bifunctional GTase/TPase) PBP2, the class B (TPase only) PBP1 and PBP3, of which PBP1 is septum-specific and has an essential function in cell division, and the low-molecular-weight PBP4 with TPase activity ^14, 16–19^. Additionally, it contains two GTases from the shape, elongation, division, and sporulation (SEDS) protein family ^9, 18, 20^, FtsW and RodA, that form cognate pairs with PBP1 and PBP3, and direct septal and lateral PG synthesis, respectively ^18^.

The essential septal PG synthases FtsW/PBP1 are thought to be activated by a trimeric complex of the divisome proteins DivIB, DivIC and FtsL (named FtsQ, FtsB and FtsL in Gram-negative bacteria, respectively), proteins conserved in cell wall-producing bacteria ^21, 22^. In *S. aureus*, the PG-binding protein DivIB is essential for septal completion and arrives at the divisome approximately at the same time as PBP1 and FtsW ^14, 23^, while DivIC spatially regulates PG synthesis by influencing the recruitment of FtsW to the division septum ^24^. We have previously shown that the *S. aureus* DivIB-DivIC-FtsL sub-complex mediates a crucial step in divisome assembly, as it is required to recruit MurJ, the flippase of the PG precursor lipid II, to the division site, driving PG incorporation to the mid-cell ^14^. Arrival of MurJ marks the transition between two stages of cytokinesis, as FtsZ treadmilling is essential for cell division at initiation of constriction before MurJ arrival, but becomes dispensable for septum synthesis in *S. aureus* afterwards ^14^.

Recent studies have indicated differences between bacterial species regarding the role of FtsZ treadmilling in cell division and the spatiotemporal distribution of PG synthases. In *Escherichia coli*, FtsZ treadmilling directs the processive movement of a fast subpopulation of non-active septal cell wall synthesis enzymes, presumably to ensure their homogeneous distribution around the division site, but does not limit the rate of septum closure ^4, 25^. However, active PG synthases move at a slower speed, independently of FtsZ treadmilling, suggesting a two-track model for active (following septal PG) and inactive (following the Z-ring) PG synthases in *E. coli* ^25^. In contrast, the rate of FtsZ treadmilling in *Bacillus subtilis* correlates both with the movement speed of septal PG synthesis enzymes and the rate of cell division ^3^. FtsZ treadmilling in this organism is essential for guiding septal cell wall synthesis during constriction initiation, after which it becomes dispensable for division while maintaining a role in accelerating the septum constriction rate ^5^. On the other hand, in *Streptococcus pneumoniae*, neither the rate of PG synthesis nor the movement of the septal PG synthesis enzyme complex are controlled by treadmilling FtsZ filaments ^26^. Despite sharing overlapping trends, these models leave open the question about the conservation of mechanisms controlling septal PG synthesis in bacteria.

Here, we studied the movement dynamics of key proteins involved in Z-ring assembly and septal PG synthesis during cell division in *S. aureus*. Using single-molecule tracking, we describe the directional movement of FtsW, PBP1 and DivIB, each existing in a single motile population, around the division site and show that their velocities are independent of FtsZ treadmilling speeds. Furthermore, we show that both FtsW GTase and PBP1 TPase activities are required for the directional movement of the septal PG synthases and that FtsW/PBP1 activity represents a rate-limiting step in cell division. We propose a one-track model for septal PG synthases in *S. aureus* featuring FtsZ treadmilling-independent movement of active FtsW/PBP1 complexes, in association with DivIB, to drive the synthesis of the new septum.

## Results

### Septal PG synthesis and not FtsZ treadmilling is rate-limiting for cell constriction

We have previously shown that cells of the coccoid bacterium *S. aureus* treated with the FtsZ inhibitor PC190723 fail to initiate the synthesis of a new division septum but do not stop ongoing constriction of existing septa, suggesting that FtsZ treadmilling is required only at the early stage of cytokinesis ^14^. Similarly, FtsZ treadmilling in rod-shaped *B. subtilis* is essential for constriction initiation, but becomes dispensable after the arrival of PG synthesis machinery, although it remains rate-limiting for cell division ^5^. We now asked whether the rate of septum constriction is accelerated by FtsZ treadmilling in *S. aureus* as well. To address this question, we constructed mutants with slowed FtsZ treadmilling speed in methicillin-resistant *S. aureus* strains COL and JE2 by replacing their native *ftsZ* gene with an allele encoding the GTPase point mutation T111A. The same amino acid substitution in FtsZ from *B. subtilis* was previously shown to cause a ∼10-fold reduction in GTPase activity without affecting the polymerization of the protein *in vitro* ^27^. The thermosensitive FtsZ(T111A) mutant derivative strains of COL and JE2 produced the FtsZ protein at wild-type levels (Supplementary Fig. 1a) and incorporated fluorescent D-amino acid HCC-amino-d-alanine (HADA) into septal PG (Supplementary Fig. 1b), despite showing heterogeneous cell size (Supplementary Fig. 2a,c) and growth rates reduced by ∼6-16% (Supplementary Fig. 3a). The FtsZ(T111A) mutation in the COL strain did not cause a delay in cell cycle progression for cells grown at 30°C, as assessed by the quantification of the frequency of cells in each cell cycle stage (Supplementary Fig. 2e). Introducing the FtsZ(T111A) mutation into the JE2 strain, and growing it at 37°C to exacerbate cell morphology defects, produced cells with multiple and misplaced septa (Supplementary Fig. 4).

We determined FtsZ treadmilling speed in nascent Z-rings using a functional fusion of EzrA, a direct interaction partner of cytoskeletal protein FtsZ, to superfast green fluorescent protein (sGFP). We used EzrA-sGFP as a proxy for FtsZ treadmilling based on our previous finding that both proteins show the same movement dynamics, sensitive to FtsZ-targeting PC190723 ^14^. As expected, FtsZ treadmilling speeds in the FtsZ(T111A) mutant strains were severely reduced relative to the parental strains (Fig. 1a,c; Supplementary Table 1; Supplementary Movie 1). To measure the rate of septum constriction in dividing COL and JE2 *S. aureus* cells, we labelled the divisome using EzrA-sGFP. Surprisingly, despite nearly complete inhibition of FtsZ treadmilling and the presence of cells with spiralling septa (Supplementary Movie 2), septum constriction rates of the FtsZ(T111A) mutants were remarkably similar to those of the respective parental strains (Fig. 1a,c; Supplementary Table 1; Supplementary Movie 2). Likewise, treating COL cells with PC190723 to inhibit FtsZ treadmilling did not significantly affect the septum constriction rate (Fig. 1b,d; Supplementary Table 1; Supplementary Movies 3,4). These results indicated that, unlike in *B. subtilis*, GTPase-driven FtsZ treadmilling is not rate-limiting for septum constriction in *S. aureus*.

**Figure 1.**
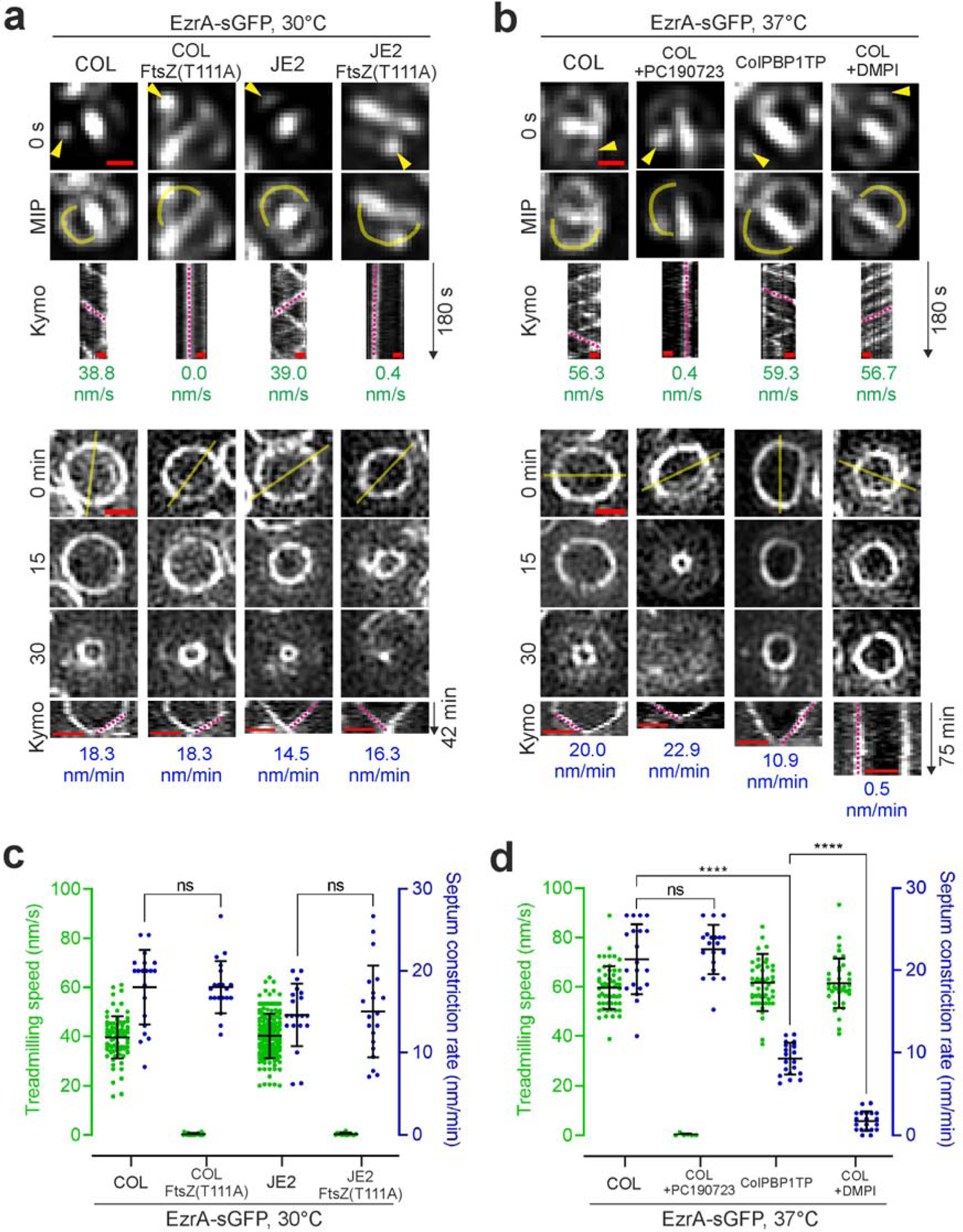
Septum constriction rate is slowed in cells lacking PBP1 TPase activity but not in FtsZ GTPase mutants impaired in treadmilling. **a,b**, Representative fluorescence micrographs of EzrA-sGFP in COL and JE2 backgrounds producing either FtsZ wild-type or thermosensitive FtsZ(T111A) mutant variants at 30°C, or in PBP1 TPase mutant ColPBP1TP and COL backgrounds at 37°C and in the absence and presence of FtsZ inhibitor PC190723 or MurJ inhibitor DMPI. The top panel shows epifluorescence images of EzrA-sGFP in late pre-divisional cells with nascent Z-rings at the start (0 s) and throughout a 180-s time series (MIP, maximum intensity projection). Yellow arrow heads indicate EzrA-sGFP patches whose changes in localization was followed over time. Space-time kymographs were generated by extracting fluorescence intensity values along indicated yellow freehand lines. Magenta dashed straight lines in kymographs indicate the slopes used to calculate EzrA-sGFP movement in nm/s, a proxy for FtsZ treadmilling speed. The bottom panel shows structured illumination images of constricting EzrA-sGFP rings acquired at indicated time points. Magenta dashed straight lines in kymographs indicate the slopes used to calculate septum constriction rates in nm/min. Scale bars, 0.5 µm. c,d, Scattered dot plots of EzrA-sGFP movement speed and ring constriction rates measured in indicated genetic backgrounds and growth conditions. c, Bars indicate the means and standard deviations of slopes obtained from lines drawn on kymographs of COL EzrA-sGFP (n_1_=83; n_2_=20), COL EzrA-sGFP FtsZ(T111A) (n_1_=21; n_2_=20), JE2 EzrA-sGFP (n_1_=171; n_2_=20;), and JE2 EzrA-sGFP FtsZ(T111A) (n_1_=15; n_2_=20). d, Bars indicate the means and standard deviations of slopes obtained from lines drawn on kymographs of COL EzrA-sGFP (n_1_=55; n_2_=20), COL EzrA-sGFP +PC190723 (n_1_=6; n_2_=20), ColPBP1TP EzrA-sGFP (n_1_=51; n_2_=20), and COL EzrA-sGFP +DMPI (n_1_=37; n_2_=20). n_1_, number of analysed slopes obtained from EzrA-sGFP movement kymographs. n_2_, number of analysed slopes obtained from EzrA-sGFP ring constriction kymographs. Statistical analysis was performed using a two-tailed Mann–Whitney *U*-test. ****, *p*<0.0001. ns, not significant.

Next, we analysed a mutant in the methicillin-resistant *S. aureus* strain COL lacking TPase activity of the division-specific PG synthase PBP1 (ColPBP1TP) ^18^. The growth rate of this mutant remains unaffected (Supplementary Fig. 3b), despite having decreased PG cross-linking ^18^, but septum constriction rate was reduced ∼2-fold (Fig. 1b,d; Supplementary Table 1; Supplementary Movie 4). In agreement, quantitative cell cycle analysis revealed a larger fraction of cells with a constricting septum for the ColPBP1TP strain relative to the parental COL strain (Supplementary Fig. 2b,d,f). This further suggested that cells lacking PBP1 TPase activity have a reduced septum synthesis rate, in line with a previous report showing a septum completion defect of COL cells depleted for PBP1 ^16^. Furthermore, treatment of COL wild-type cells with DMPI, an inhibitor of the lipid II flippase MurJ that blocks PG synthesis, almost completely halted septum constriction (Fig. 1b,d; Supplementary Table 1; Supplementary Movie 4), in accordance with our previous data ^14^. Importantly, in both conditions where the septum constriction rate was slowed down, FtsZ treadmilling speed remained unchanged (Fig. 1b,d; Supplementary Table 1; Supplementary Movie 3). These results suggest that septal PG synthesis resulting from FtsW/PBP1 activity, not FtsZ treadmilling, is the primary driver of cell constriction in *S. aureus*.

### FtsW, PBP1 and DivIB move directionally around the division site

Our data indicate that PBP1 TPase activity is a major driver of septum constriction. In *S. aureus*, PBP1 acts in concert with its cognate SEDS family protein FtsW ^18^. The homolog division SEDS-bPBP pairs in *B. subtilis, S. pneumoniae*, *E. coli* and *Caulobacter crescentus* undergo directional movement at the division site ^6,25, 26, 28^. We aimed to investigate whether FtsW/PBP1, and other components of the cell division machinery, show similar movement behaviour in cells of methicillin-resistant *S. aureus*. To this end, we constructed self-labelling Halo-tag (HT) ^29^ and Snap-tag (ST) ^30^ fusions to various cell division and cell wall-related proteins in the background of strain JE2 EzrA-sGFP. Genomic *ftsW*, *murJ*, *pbp4* and *gpsB* were replaced with gene derivatives encoding C-terminal translational fusions to HT. Strains expressing *ht* fused to *ftsW* and *murJ* genes from the native locus, under the control of their native promoter, as the sole copy of the gene in the cell, were viable, indicating functionality of the corresponding protein fusions, given that both genes are essential in methicillin-resistant *S. aureus* ^14^. Expression of gene fusions inserted at the ectopic *spa* locus was driven either by the isopropyl β-d-1-thiogalactopyranoside (IPTG)-inducible *spac* promoter (*ht-divIB* and *rodA-ht*) or by the anhydrotetracycline (Atc)-inducible *xyl-tetO* promoter (*ist-pbp1*, *ftsZ-ht* and *ezrA-ht*). Our attempts to construct functional fusions to *pbp2* and *pbp3* were unsuccessful. None of the protein fusions produced in *S. aureus* affected cell growth, except for GpsB-HT (Supplementary Figs. 5a & 6a). Moreover, HT and ST protein fusions were not proteolytically cleaved and were enriched in the septal region of dividing cells as expected (Supplementary Figs. 5b,c & 6b,c). The exception was iST-PBP1 that showed additional cytoplasmic fluorescence likely attributed to partial proteolytic cleavage of the protein fusion (Supplementary Fig. 6b,c). We then studied by single-molecule tracking microscopy the movement dynamics at the division site, which was labelled with early cell division protein EzrA fused to sGFP, of the nine functional HT and ST fusions to various cell division and cell wall-related proteins. To resolve single molecules of HT and ST protein fusions, exponentially growing cells were sparsely labelled with JF549-HTL and JFX650-STL, respectively, and imaged by time-lapse fluorescence microscopy. Directional movement around the division site was observed for single molecules of FtsW-HT, iST-PBP1, and HT-DivIB, in both early and late stages of septum constriction (Fig. 2, Supplementary Table 2; Supplementary Movie 5). The travelling distance of molecules could span the entire septum circumference during the 180 s observation period, corresponding to a track length of ∼3 μm, and a fraction of molecules transitioned between clockwise and counter-clockwise circumferential movement (Fig. 2). Interestingly, directional movement of HT-DivIB was lost upon deletion of its C-terminal γ domain (residues 373 to 439 of DivIB) (Supplementary Table 2), which largely retained the fusions’ stability but caused mis-localization (Supplementary Fig. 7a,e). No directional movement around the division site was observed for single molecules of MurJ-HT, PBP4-HT, GpsB-HT, RodA-HT, FtsZ-HT, and EzrA-HT (Supplementary Table 2).

**Figure 2.**
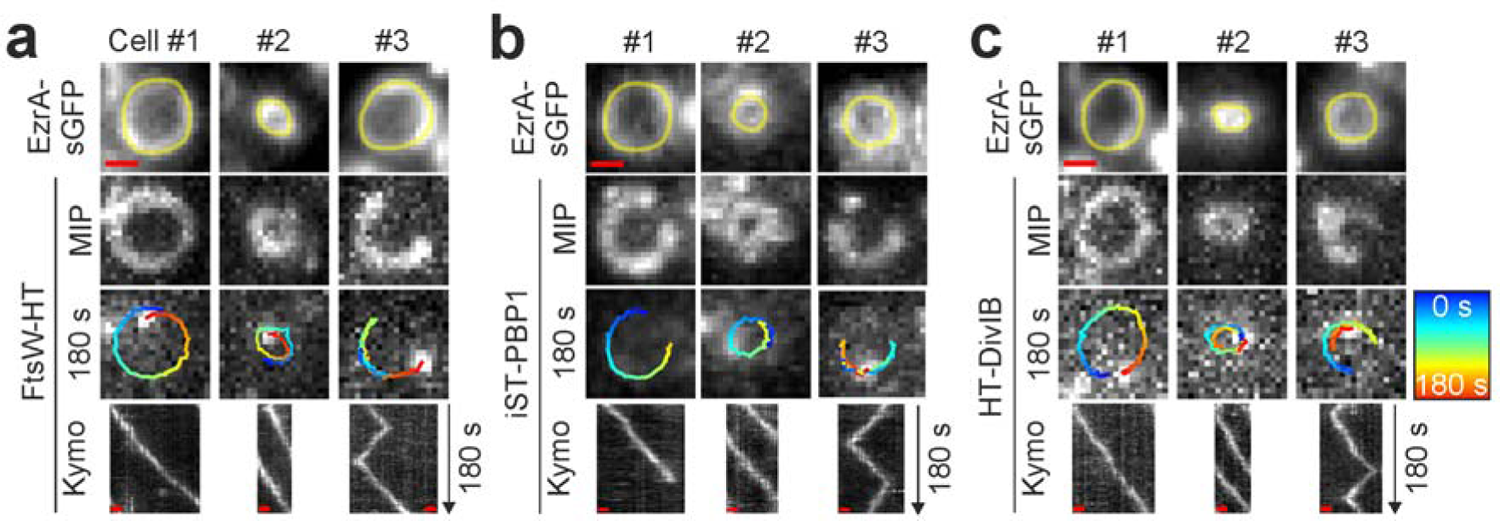
FtsW, PBP1 and DivIB move directionally around the division site. **a-c**, Representative epifluorescence micrographs of JE2 EzrA-sGFP background producing FtsW-HT (**a**), iST-PBP1 (**b**), or HT-DivIB (**c**), in TSB rich medium at 37°C. Cells in panel **a** produced FtsW-HT from its native genomic locus as the sole source of FtsW. Cells producing iST-PBP1 or HT-DivIB were grown in the presence of 2 ng/ml Atc or 0.5 mM IPTG, respectively, to induce gene expression from the ectopic *spa* locus. Cells were sparsely labelled with the fluorescent ligands JF549-HTL or JFX650-STL to visualize single molecules of FtsW-HT and HT-DivIB or iST-PBP1, respectively. Each panel shows three independent cells, of which #1 and #2 are at an early and a late stage of septum constriction, respectively, and #3 exhibits a labelled molecule transitioning between clockwise and counter-clockwise movement around the division site. Single-molecule images acquired in the last frame of a 180-s time series are overlayed with tracks, where blue (0 s) to red (180 s) indicates trajectory time. Space-time kymographs were generated by extracting fluorescence intensity values from FtsW-HT, iST-PBP1 and HT-DivIB images along yellow freehand lines drawn on corresponding EzrA-sGFP rings acquired in the last frame of each time series. MIP, maximum intensity projection of all 61 time frames. Scale bars, 0.5 µm.

The movement of these molecules was not analysed further due to their lack of continuous tracks with at least 30 data points in the selected imaging conditions and above the cut-off for track directionality (α_MSD_ ≥1) (Supplementary Table 2). We concluded that divisome proteins have distinct movement dynamics at the division site, with FtsW, PBP1 and DivIB moving directionally.

### FtsW, PBP1 and DivIB move with the same velocity

To characterise the directional movement of FtsW, PBP1 and DivIB molecules, we started by determining their velocity. *S. aureus* cells are approximately spherical and, when placed on a microscope slide, will present the division plane in random orientations. Therefore, our analysis considered single-molecule motion in all three spatial dimensions by making two assumptions: (i) directionally moving molecules moved on division rings, and (ii) the cytokinetic ring has a near-circular shape. Briefly, the molecules’ Z-position (position along the axis perpendicular to the imaging plane) was inferred by aligning their X-Y position with the nearest point in an ellipse manually drawn on a corresponding EzrA-sGFP ring, and rotating this ellipse by Θ (angle of the division ring relative to the imaging plane) into a circle (Fig 3a; Supplementary Movie 6). In order to estimate molecular velocity, we analytically unwrapped each track into a 1D representation. Tracks were then sectioned using breakpoint detection, velocity calculation was performed based on simple linear regression for each section, and the average velocity of the sections of each track was calculated (Fig. 3b). Following this approach, consistent molecule velocities were obtained over a range of Θ from 0 to ∼75°. We noticed that tracks in cells with a division plane nearly perpendicular (Θ>75°) to the imaging plane resulted in higher velocities and larger standard deviations (Supplementary Fig. 8a-c) due to frequent unwrapping artifacts, and for this reason, only tracks with Θ<75° were included in further analysis.

**Figure 3.**
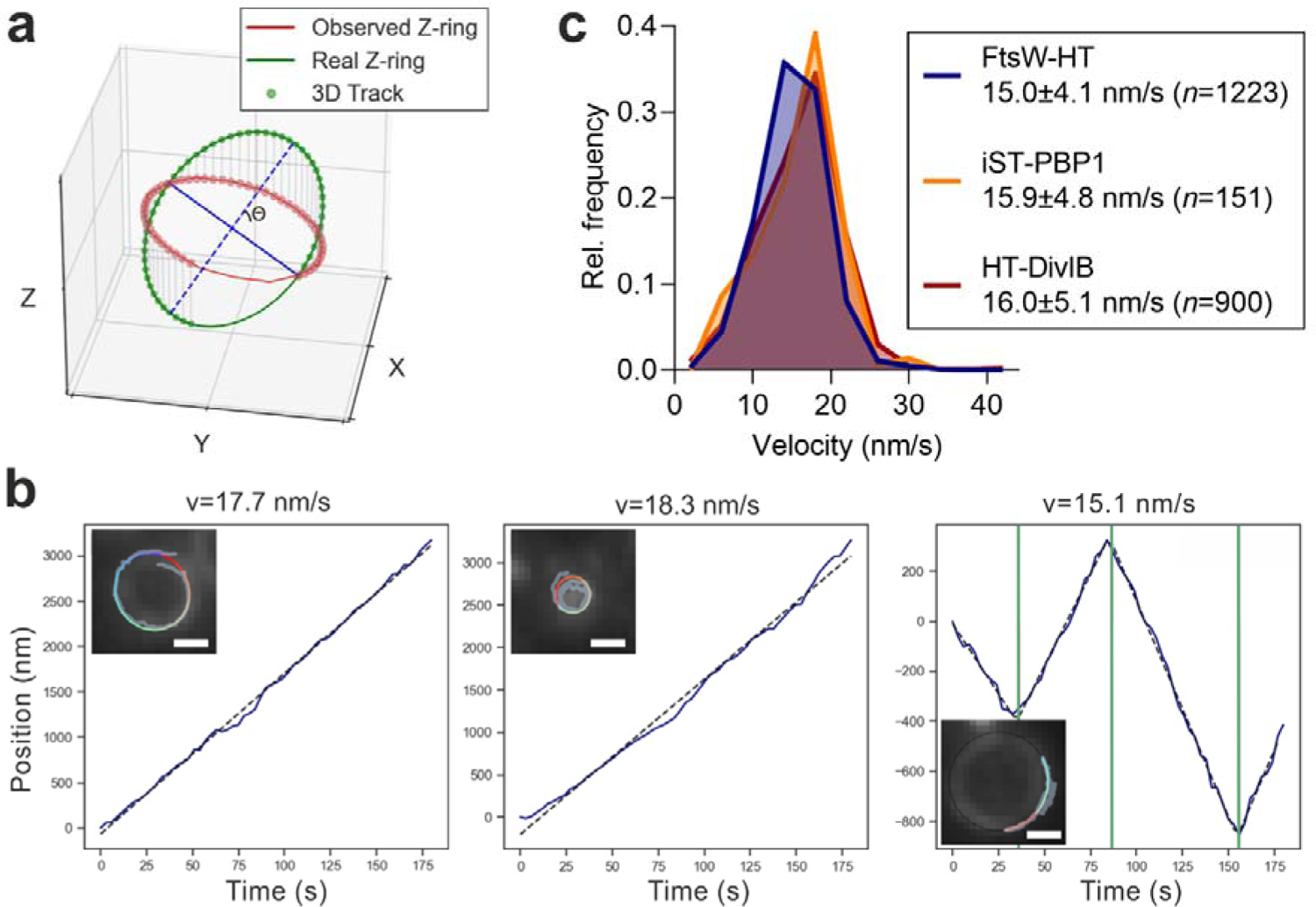
FtsW, PBP1 and DivIB move directionally with the same velocity. **a**, Representation of a simulated single-molecule track of FtsW-HT aligned to an ellipse that was drawn on top of an observed EzrA-sGFP ring. Θ indicates the angle between the imaging and division planes. **b,** Examples of unwrapped FtsW-HT trajectories (blue lines in graphs). Left and middle graphs show FtsW-HT in cells at an early and a late stage of septum constriction, respectively, and the right graph shows FtsW-HT transitioning between different directions. Green vertical lines indicate detected breakpoints used to segment tracks to perform a linear regression in each section (indicated by black dashed lines). Average velocity (v) was calculated as the mean of velocities determined for each section. Insets show corresponding micrographs of EzrA-sGFP overlayed with the original track (blue) and the same track after alignment with an ellipse (blue to red). Scale bars, 0.5 µm. **c,** Histograms of FtsW-HT, iST-PBP1 and HT-DivIB single-molecule velocities determined in JE2 EzrA-sGFP background in TSB rich medium at 37°C. Average velocity is shown as mean with standard deviation. Bin width, 4. Center of first/last bin, 2/42. Histograms were obtained from at least six biological replicates.

In JE2 and COL strains, the velocities of FtsW-HT, iST-PBP1, and HT-DivIB, directionally moving for at least 87 s (equivalent to 30 data points in a track), were similar (∼15-16 nm/s) (Fig. 3c; Supplementary Table 1). Moreover, we found no correlation between their velocity and either track duration or cell division stage (Supplementary Fig. 8d-i). The similar velocities observed for FtsW, PBP1 and DivIB throughout all stages of constriction suggest that these proteins exist in a complex which moves at a constant rate from initiation to completion of septum constriction.

### FtsW/PBP1 and DivIB velocity correlates with cell growth rate

Having established a quantitative approach to determine the average velocity of a directionally moving molecule, we used it to analyse FtsW and DivIB trajectories measured in fast and slow growth conditions. From here on, we used FtsW velocity as a proxy for that of PBP1 because both proteins were previously shown to directly interact at the division site ^18^. We found that FtsW-HT and HT-DivIB average velocities in cells of the JE2 strain grown in M9 minimal medium were reduced ∼1.3-fold relative to cells grown in TSB rich medium (Fig. 4a,b; Supplementary Table 3). FtsW and DivIB average velocities in TSB rich medium were reduced ∼1.6-fold when the growth temperature was reduced from 37°C to 30°C, and ∼2.3-fold when the temperature was decreased from 37°C to 25°C (Fig. 4a,b; Supplementary Table 3). As expected, cell growth rate was lower in M9 minimal medium or at lower temperatures than in TSB rich medium at 37°C (Fig. 4c). Plotting FtsW-HT and HT-DivIB average velocity as a function of cell growth rate indicated a positive linear correlation between these two parameters (Fig. 4d). This correlation may suggest that the directional movement of FtsW and DivIB underlies a process that is limiting for cell growth or that growth rate is limited by a factor that also restricts FtsW and DivIB velocity.

**Figure 4.**
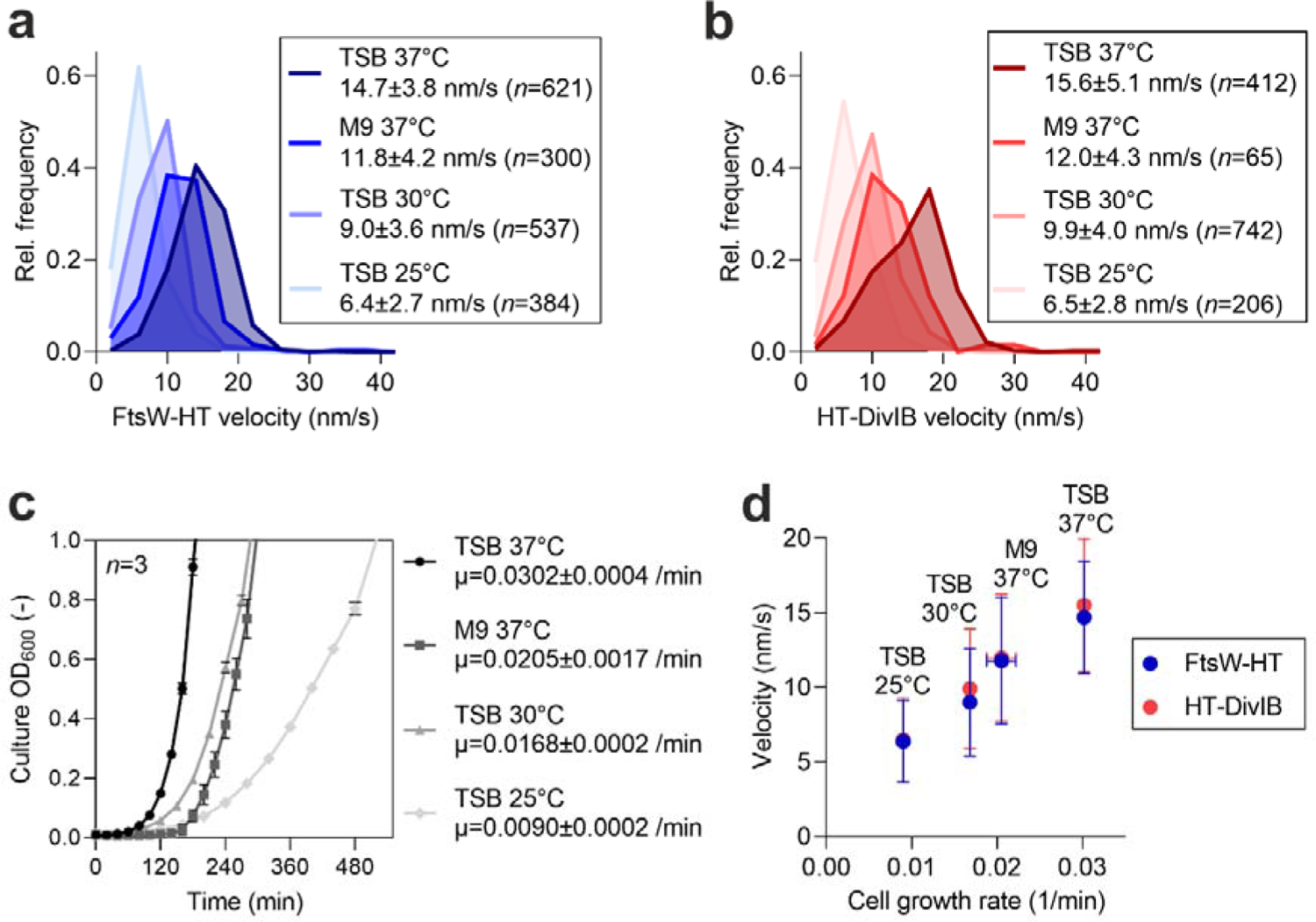
FtsW and DivIB velocity correlates with cell growth rate. **a,b**, Histograms of FtsW-HT (**a**) and HT-DivIB (**b**) single-molecule velocities determined in JE2 EzrA-sGFP background, in rich and poor media and at various growth temperatures. Average velocity is shown as mean with standard deviation. Bin width, 4. Centre of first/last bin, 2/42. **c,** Mean optical density of culture as a function of time determined for strain JE2 EzrA-sGFP grown in indicated media and at various temperatures. Error bars represent the standard deviations of three biological replicates. **d,** FtsW-HT and HT-DivIB mean velocities shown in panels **a** and **b** as a function of cell growth rate determined from growth curves shown in panel **c**. Horizontal error bars represent the standard deviations from three biological replicates and vertical error bars represent the standard deviations of a minimum of 65 trajectories from three biological replicates.

### Directional movement of FtsW or DivIB does not depend on FtsZ treadmilling

Previous studies using different bacterial models have shown the directional movement of septum-specific PG synthases to be dependent or independent of treadmilling FtsZ filaments, depending on the species ^3,4, 25, 26^. To examine whether FtsW and DivIB velocities are correlated with FtsZ treadmilling speed in *S. aureus*, we first studied the effect of impaired FtsZ treadmilling on the directional movement of FtsW and DivIB using FtsZ(T111A) mutants. This mutation, which did not affect FtsW-HT co-localization with EzrA-sGFP at the division site (Supplementary Fig. 1b), severely reduced FtsZ treadmilling speeds in both JE2 and COL backgrounds (Fig. 1a,c; Supplementary Table 1; Supplementary Movie 1). Despite this dramatic decrease, FtsW-HT and HT-DivIB velocities were not markedly changed (Fig. 5a,b; Supplementary Tables 1,3; Supplementary Movie 7). Similarly, PC190723 treatment of cells, which virtually stopped FtsZ treadmilling (Fig. 1b,d; Supplementary Table 1; Supplementary Movie 3), did not affect FtsW-HT or HT-DivIB velocities (Fig. 5c,d; Supplementary Tables 1,3; Supplementary Movie 5). Importantly, FtsW-HT and HT-DivIB velocities were not affected by the FtsZ(T111A) mutation or PC190723 treatment at any stage of cell division (Supplementary Fig. 9). Moreover, FtsW-HT and HT-DivIB velocities in JE2 and COL backgrounds, at 30°C and 37°C, were ∼4-fold slower than FtsZ treadmilling speeds at the respective growth temperature (Supplementary Table 1), indicating that FtsW and DivIB do not move together with FtsZ. Combined, these results suggest that the movement of FtsW and DivIB along septal rings in *S. aureus* is not coupled to the treadmilling of FtsZ filaments.

**Figure 5.**
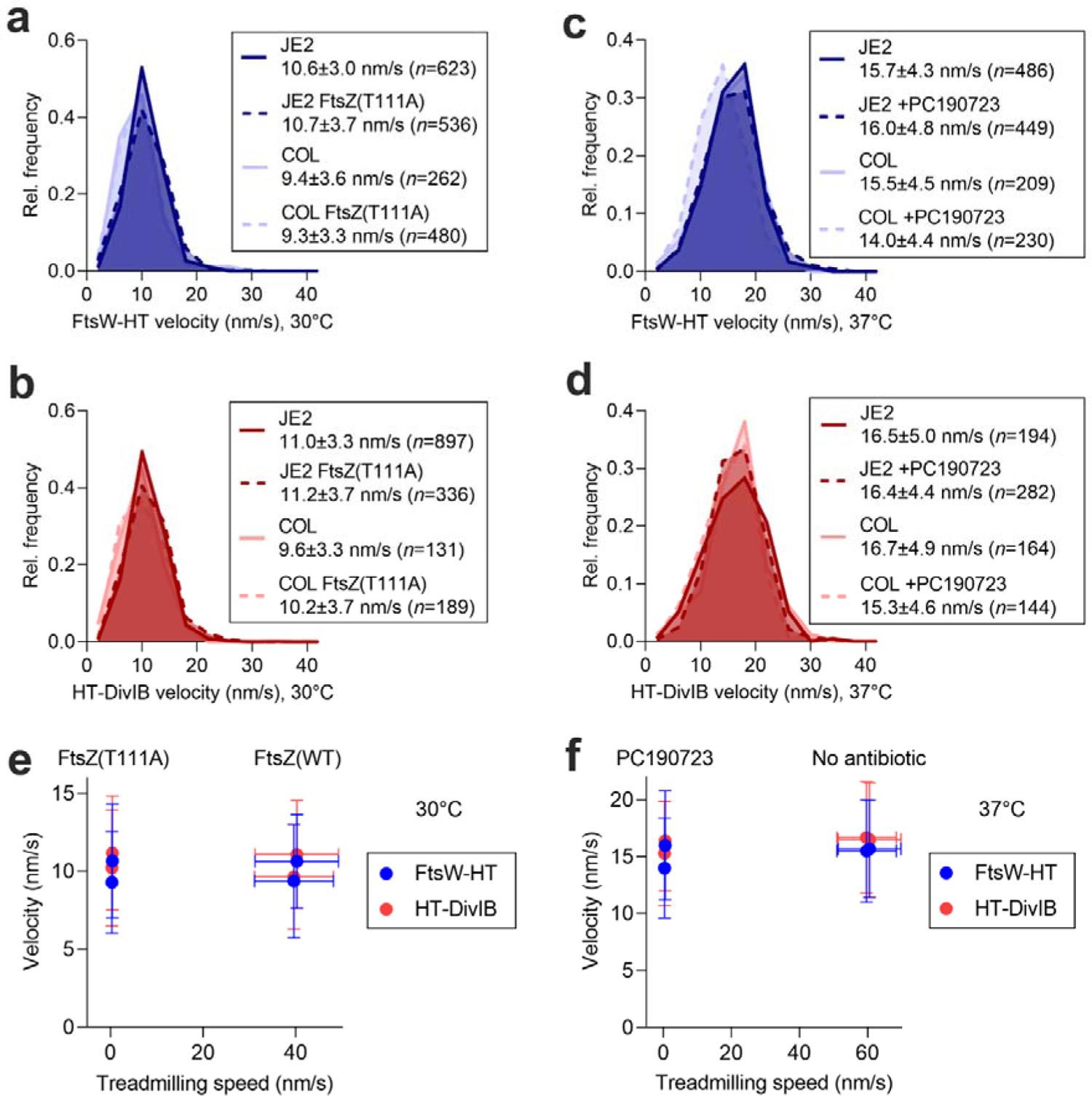
FtsW and DivIB velocity does not correlate with FtsZ treadmilling speed. **a-d**, Histograms of FtsW-HT and HT-DivIB single-molecule velocities determined in JE2 EzrA-sGFP and COL EzrA-sGFP backgrounds, each producing either FtsZ wild-type or temperature-sensitive FtsZ GTPase mutant T111A from its native genomic locus in TSB rich medium at 30°C (**a,b**), or in the absence and presence of FtsZ inhibitor PC190723 in TSB rich medium at 37°C (**c,d**). Average velocity is shown as mean with standard deviation. Bin width, 4. Center of first/last bin, 2/42. **e,f,** FtsW-HT and HT-DivIB mean velocities shown in panels **a-d** as a function of FtsZ treadmilling speed shown in Figure 1 and Supplementary Table 1. Horizontal error bars represent the standard deviations of a minimum of 6 slopes and vertical error bars represent the standard deviations of a minimum of 131 trajectories from three biological replicates.

### PG synthesis by FtsW/PBP1 drives the directional movement of FtsW/PBP1 and DivIB

Given that FtsZ treadmilling was not the driver for the directional movement of FtsW/PBP1 and DivIB in *S. aureus*, we next asked whether their velocity was determined by PG synthesis. Addition to cells of DMPI, an inhibitor of the lipid II flippase MurJ, did not slow down FtsZ treadmilling speed (Fig. 1b,d; Supplementary Table 1; Supplementary Movie 3), but caused a ∼3-fold reduction in FtsW-HT and HT-DivIB velocities (Fig. 6a,b; Supplementary Tables 1,3; Supplementary Movie 5). Treatment of cells with imipenem, a β-lactam with preferential activity against *S. aureus* PBP1 ^31^, also resulted in a ∼2-fold reduction in FtsW-HT and HT-DivIB velocities (Fig. 6a,b; Supplementary Table 3; Supplementary Movie 5). Strikingly, treatment of cells with the glycopeptide antibiotic vancomycin that inhibits PG synthesis, previously shown to cause a total arrest of cell division at all stages in *S. aureus* within minutes ^32^, nearly completely stopped the directional movement of FtsW-HT and HT-DivIB (Supplementary Fig. 10; Supplementary Movie 5). The frequency of cells exhibiting directionally moving molecules was reduced from ∼6-8% in untreated to ∼0.1-0.2% in vancomycin-treated cells (Supplementary Table 3). Noteworthy, vancomycin treatment did not affect FtsW-HT and EzrA-sGFP localization to mid-cell, suggesting that divisomes remained intact in these cells (Supplementary Fig. 7b).

**Figure 6.**
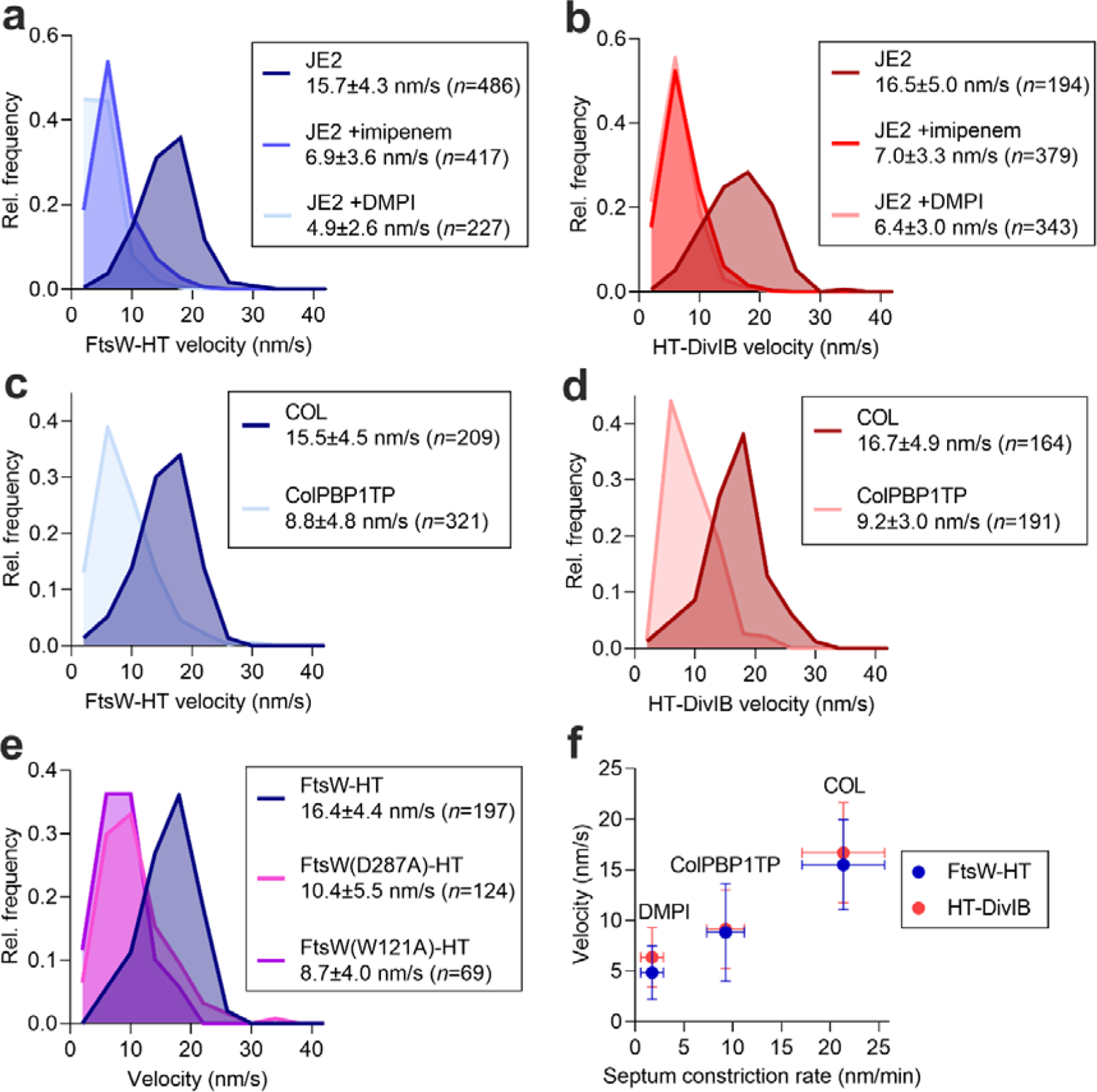
Inhibition of peptidoglycan synthesis slows down the directional movement of FtsW and DivIB. **a-e**, Histograms of FtsW-HT and HT-DivIB single-molecule velocities determined in JE2 EzrA-sGFP and COL EzrA-sGFP backgrounds in the absence and presence of beta-lactam imipenem or MurJ inhibitor DMPI (**a,b**), in PBP1 TPase mutant ColPBP1TP EzrA-sGFP and COL EzrA-sGFP backgrounds at 37°C (**c,d**), and in JE2 EzrA-sGFP background producing either FtsW-HT wild-type or its active-site mutant derivatives W121A and D287A from the ectopic *spa* locus (**e**). Average velocity is shown as mean with standard deviation. Bin width, 4. Center of first/last bin, 2/42. **f,** FtsW-HT and HT-DivIB mean velocities shown in panels **a-d** as a function of septum constriction rate shown in Figure 1 and Supplementary Table 1. Horizontal error bars represent the standard deviations of 20 cells and vertical error bars represent the standard deviations of a minimum of 164 trajectories from three biological replicates.

We then used active-site mutants of both proteins to test which PG synthesis activity, FtsW GTase or PBP1 TPase, was required for the movement of FtsW and DivIB. Inactivation of PBP1 TPase activity via a S314A mutation in the ColPBP1TP strain, decreased FtsW-HT and HT-DivIB velocities by ∼1.5-2-fold relative to the parental COL strain (Fig. 6c,d; Supplementary Table 3; Supplementary Movie 8), while track duration and FtsW-HT localization to mid-cell remained unchanged (Supplementary Fig. 7c; Supplementary Table 3). Next, the GTase point mutations W121A and D287A, previously shown to impair the essential function of FtsW in *S. aureus* ^18^, were introduced into the FtsW-HT fusion produced from the ectopic *spa* locus. As a control, we also introduced wild-type FtsW-HT in the *spa* locus and confirmed that its velocity was the same when produced either from the ectopic *spa* locus or from its native locus (Fig. 6a,e; Supplementary Table 3). Relative to ectopic FtsW-HT, the velocities of the FtsW GTase mutants were reduced between ∼1.5- and ∼2-fold, while all protein variants localized at mid-cell and were subject to no apparent proteolytic cleavage (Fig. 6e; Supplementary Fig. 7d,e). Slowed velocities of FtsW GTase mutant variants were accompanied by reductions in the percentage of cells with tracks from ∼3.8% to ∼0.4-0.9% (Supplementary Table 3). These results indicate that the processive movement of FtsW/PBP1 complexes and DivIB along septal rings is driven by FtsW GTase and/or PBP1 TPase enzyme activities.

### FtsW and DivIB exist in a single motile population

Previous studies using Gram-negative *E. coli* and *C. crescentus* proposed a two-track model where directionally moving FtsW exists in two motile populations, one slow (active) depending on PG synthesis and one fast (inactive) depending on FtsZ treadmilling ^25, 28^. To see if this was also the case for *S. aureus*, we used the same analysis as above but without averaging the velocities of each track. This allows us to detect cases where molecules would switch between slow and fast movement, giving rise to bimodal distributions. Through this analysis we verified that FtsW-HT velocities show a unimodal distribution (Supplementary Fig. S11). In *E. coli*, depletion of the slow (active) subpopulation of FtsW through inhibition of PG synthesis, leads to an increase of the overall average speed of FtsW molecules^25^. We did not observe a shift towards higher velocities, nor an appearance of a new fast population after decreasing the number of processively moving FtsW-HT molecules in *S. aureus*, either by slowing down the cell growth rate, inhibiting PG synthesis by DMPI treatment, or inactivating PBP1 TPase and FtsW GTase (Supplementary Fig. S11c,e,g). The velocity distribution of HT-DivIB showed the same unimodal distribution in these conditions (Supplementary Fig. S11d,f). To rule out the possibility that our imaging conditions did not allow for the detection of FtsW moving with a velocity that would scale with FtsZ treadmilling speed in fast-growing cells (∼60 nm/s), we determined the FtsW-HT velocity distribution using three times faster image acquisition rate (increase from 0.33 Hz to 1 Hz). However, similar FtsW-HT velocity distributions, showing the unimodal distribution, were still observed (Supplementary Fig. S11h). Combined with our finding that inhibition of PG synthesis by vancomycin virtually stopped the directional movement of FtsW-HT and HT-DivIB (Supplementary Fig. 10; Supplementary Table 3; Supplementary Movie 5), this data suggests these two proteins exist in a single motile population depending on PG synthesis.

### FtsW and DivIB velocity correlates with septum constriction rate

Here, we identified PBP1 TPase and FtsW GTase activities determining the velocity of directionally moving FtsW (Fig. 6c,e), and PBP1 TPase activity to be rate-limiting for septum constriction (Fig. 1b,d). PG synthesis is likely the sole driver for the directional movement of FtsW and DivIB (both were virtually stopped by vancomycin; Supplementary Fig. 10; Supplementary Table 3; Supplementary Movie 5), and therefore their velocities serve as an indicator for the rate of septal PG synthesis. Previous data have shown that the cell division process in *S. aureus* depends on PG synthesis at all stages, as its inhibition by vancomycin is sufficient for a total arrest of septum constriction ^32^. Based on these observations, we hypothesized that the rates of septal PG synthesis and septum constriction may correlate. Plotting the velocities of FtsW-HT and HT-DivIB as a function of septum constriction rate, using data points obtained for COL, the ColPBP1TP mutant lacking PBP1 TPase activity, and cells treated with MurJ inhibitor DMPI, revealed a positive linear correlation between these two parameters (Fig. 6f). This data supports a model in which the enzymatic activities by FtsW/PBP1 complexes, in concert with DivIB, determine the rate of septal PG synthesis and hence the rate of septum synthesis in *S. aureus* (Fig. 7).

**Figure 7.**
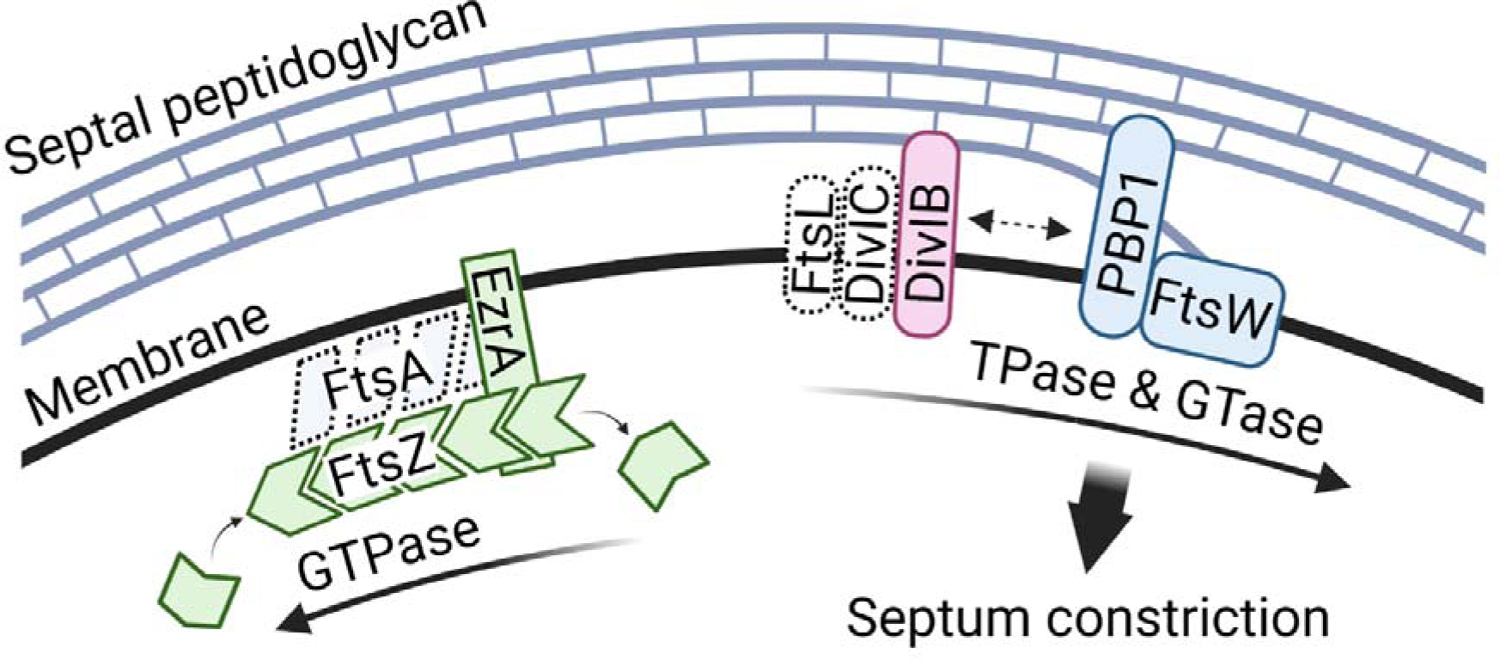
Model for the movement dynamics of essential cell division proteins in *S. aureus*. The directional movement of the septal peptidoglycan synthases FtsW and PBP1 is driven by both their glycosyltransferase (GTase) and transpeptidase (TPase) activities, respectively. Processively moving FtsW/PBP1 remains associated with the activating sub-complex DivIB-DivIC-FtsL to synthesize new peptidoglycan and thereby promote inward growth of the division septum. In this model, GTPase-driven treadmilling by FtsZ filaments neither determines the velocity of the essential PG synthesis enzyme complex nor the rate of septum synthesis in *S. aureus*. Note that FtsA, DivIC and FtsL movement dynamics were not analysed in this study.

## Discussion

In most microscopy studies on cytokinesis of different ovococci and rod-shaped bacterial models, cells are imaged with their division plane oriented vertically relative to the imaging plane, posing a limitation to imaging depth and spatial resolution ^4,6, 25, 28^. This limitation was recently overcome by imaging vertically immobilized bacteria, where the division plane is parallel to the imaging plane ^3, 5, 26, 33^. This laborious methodology enabled more accurate measurement of the motion of cell division proteins, but the collection of large data sets has remained a challenge, especially for single-molecule imaging ^33^. In contrast to rods and ovococci, spherical cells spotted on a microscope slide lay in all possible orientations and often have their division plane oriented parallel to the imaging plane; this allows imaging with high spatial resolution and facilitates vast data collection to study the mechanisms underlying cytokinesis. We took advantage of this fact and used time-resolved fluorescence microscopy to examine the movement dynamics of cell division and cell wall-related proteins during cytokinesis in the coccoid bacterium *S. aureus*.

We analysed the movement dynamics of cell division proteins arriving early (FtsZ, EzrA) or late (FtsW, PBP1, DivIB, GpsB, MurJ) at the division site, as well as of the elongasome protein RodA and PG TPase PBP4. Single-molecule imaging robustly detected directional movement around the division plane only for FtsW, PBP1 and DivIB. Consistent with previous data, FtsZ and EzrA move around the division site in treadmilling patches/filaments rather than as single molecules ^5,6, 14^. Previous single-molecule imaging of *B. subtilis* revealed two dynamically distinct sub-complexes of the divisome, one composed of stationary FtsZ binding proteins (FtsA, EzrA, SepF, ZapA), and one containing directionally moving cell wall synthases (FtsW, PBP2b, DivIB, DivIC, FtsL) ^6^. Given that, in this study, FtsW/PBP1 and DivIB showed the same directional movement in all conditions tested, we hypothesize that FtsW/PBP1 engage in a stable complex with sub-complex DivIB-DivIC-FtsL in *S. aureus*. This hypothesis is in agreement with a recent study describing the isolation of the *Pseudomonas aeruginosa* septal PG synthesis enzyme complex comprising the proteins FtsQ (DivIB ortholog), FtsB (DivIC ortholog), FtsL, FtsW and FtsI (ortholog of *S. aureus* PBP1) ^34^. The *S. aureus* protein DivIB contains an N-terminal cytoplasmic domain, a transmembrane segment and an extracytoplasmic region composed of α, β and γ domains. Our data indicate that the C-terminal γ domain of *S. aureus* DivIB is important for its recruitment to mid-cell and for stabilising active FtsW/PBP1 complexes. This would be similar to *E. coli* where structure prediction and molecular dynamics simulations suggest a role of FtsQ in stabilizing the FtsQLBWI complex through its C-terminal domains ^35^. When similar methods are applied to the orthologous complex in *S. aureus*, the predicted structure of the complex (Supplementary Fig. S12) is similar to that predicted for and observed by cryo-EM for Gram-negative bacteria ^34, 35^. The DivIB β and γ domains seem to play roles in mediating the predicted interaction between DivIB and the PBP1 PASTA domains, which was reported in *B. subtilis* ^36^. We have previously shown that DivIB-DivIC-FtsL is required to recruit the putative lipid II flippase MurJ ^14^, thereby setting on septal PG incorporation. Since we were unable to robustly detect directional movement of a functional MurJ fusion, our data suggest that in *S. aureus* MurJ does not stably interact with a putative pentameric complex of FtsW, PBP1, DivIB, DivIC and FtsL.

Our findings indicate that FtsW/PBP1 velocity scales with constriction rate and remains constant at all stages of cytokinesis, suggesting that, once initiated, cell constriction in *S. aureus* is a continuous process driven by PG synthesis. In our experiments, PG synthesis was inhibited either by using active site mutants of FtsW and PBP1, or by treating cells with antibiotics targeting different steps in the PG synthesis pathway: MurJ-inhibitor DMPI, which lowers extracellular levels of the PG precursor lipid II, imipenem, which inhibits PBP enzymes, and vancomycin, which binds to PG precursor lipid II. In all cases, FtsW’s directional movement was either slowed down or completely stopped, suggesting that directionally moving FtsW/PBP1 complexes in *S. aureus* are enzymatically active (Fig. 7). Based on the strict dependence of FtsW movement on septal PG synthesis, we hypothesize that tracks spanning (almost) the entire circumference of the division site may correspond to the synthesis of long glycan strands. Similarly, it is possible that the FtsW/PBP1 observed transitioning between clockwise and counter-clockwise circumferential movement, may correspond to protein complexes initiating the synthesis of a new glycan strand in the opposite direction.

The discovery of FtsZ treadmilling behaviour prompted the question of the role of this process in bacterial cell division ^2^. One of the earlier hypotheses was that FtsZ treadmilling was required to distribute PG synthases around the division site homogeneously. This hypothesis was reinforced when direct evidence of the PG synthase movement dependency on FtsZ treadmilling was provided for *B. subtilis* and *E. coli* ^3, 4^. FtsZ GTPase mutants in *E. coli* reduced the population size of fast-moving (inactive) FtsWI complexes and, therefore, the overall velocity of directionally moving PG synthases ^4, 25^. A reduction of FtsZ treadmilling speed in *B. subtilis*, induced either by producing FtsZ GTPase mutants or by treating cells with PC190723, caused directionally moving PG synthases to slow down or become immobile ^3^. We did not observe a similar behaviour in *S. aureus.* In our experiments, when FtsZ treadmilling was impaired either by introducing the GTPase mutation T111A into the native FtsZ protein of two *S. aureus* strains or by treating cells with PC190723, the velocities of directionally moving FtsW and DivIB remained unchanged. Even at the initial stage of cell division, where FtsZ treadmilling was shown to be essential and become dispensable only at a later stage ^5, 14^, we could not find a dependency of FtsW or DivIB directional movement on FtsZ treadmilling (Supplementary Fig. S9). This result is in contrast with the studies of *E. coli* ^4^ and *B. subtilis* ^3^, but in line with a previous report on *S. pneumoniae*, in which PG synthase movement depends on PG precursor availability rather than FtsZ treadmilling ^26^. *S. pneumoniae* FtsZ GTPase mutants impaired in treadmilling did not reduce the velocities of FtsW or bPBP2x (ortholog of *S. aureus* PBP1) or reduced them only slightly ^26^. To explain the link between the directional movement of PG synthases and FtsZ treadmilling, an *in vitro* study proposed a diffusion-and-capture mechanism for divisome proteins following treadmilling FtsZ filaments ^37^. According to a Brownian ratchet model, bound FtsI molecules would undergo end-tracking at the shrinking end of the FtsZ polymer ^33^. Although our data suggest the directional movement of FtsW to be uncoupled from FtsZ treadmilling, we cannot exclude that FtsW molecules may track with FtsZ filaments for a very short time, while spending most of the time actively synthesizing PG using lipid II as substrate.

The question therefore remains regarding the essential function of FtsZ treadmilling in cell division*. E. coli* mutants with severely reduced FtsZ treadmilling were substantially altered in the ultrastructure of the septal cell wall and showed polar morphology defects ^4^. In *B. subtilis,* FtsZ treadmilling is essential for Z-ring condensation, guides the initiation of constriction and is rate-limiting for cell division ^3,5^. Specifically, the FtsZ GTPase mutation T111A caused the formation of mini-cells in *B. subtilis* and resulted in a ∼1.6-fold decrease in FtsZ treadmilling speed, accompanied by a ∼1.5-2-fold reduction in septum constriction rate ^3,27^. We now showed that an *S. aureus* strain containing the same FtsZ mutation, as well as cells treated with PC190723, share some phenotypes: FtsZ treadmilling was severely reduced, a larger fraction of cells showed constriction defects and their morphology was substantially altered. However, contrarily to *B. subtilis*, the rate of septum synthesis remained unchanged even in the presence of PC190723, which completely stops FtsZ treadmilling ^14^. Overall, we propose that, in *S. aureus*, FtsZ treadmilling does not determine the rate of septal PG synthesis but rather constitutes a mechanism for ensuring uniformity of septum structure and hence the formation of two equally sized daughter cells.

In this work we addressed one final question, regarding the existence of one or two subpopulations of processively moving PG synthases in *S. aureus.* In Gram-negative *E. coli* directionally moving FtsWI exists in two motile populations: actively PG-synthesizing complexes move slowly (∼8 nm/s) relative to their enzymatically inactive counterparts (∼30 nm/s), and only the latter scale with FtsZ treadmilling speed (∼28 nm/s) ^4,25^. Two motile populations were also observed in Gram-negative *C. crescentus* ^28^. We provide three pieces of evidence to support the conclusion that in *S. aureus* FtsW exists in a single motile population that depends on PG synthesis and not on FtsZ treadmilling: (i) FtsW moved with a slower velocity than treadmilling FtsZ filaments, (ii) FtsW velocity remained unchanged when FtsZ treadmilling was impaired, (iii) inhibition of PG synthesis slowed down or completely stopped FtsW, but not FtsZ treadmilling, and did not reveal a fast (inactive) FtsW subpopulation. Given that two populations of directionally moving FtsW were detected only in Gram-negative species so far, it is tempting to speculate that FtsW dynamics may correlate with the different amounts of PG produced by bacteria. Gram-positive PG contains many layers and is 30-100 nm thick, whereas Gram-negative PG has only one to a few layers ^38^. Thus, directionally moving FtsW/bPBP in Gram-positive bacteria could exist mostly in its active form to synthesize the large amount of septal PG required for cell division. To produce a relatively small amount of septal PG at a similar synthesis rate, FtsW in Gram-negative species may spend more time in its enzymatically inactive state, during which it could track with treadmilling FtsZ filaments.

## Methods

### Bacterial growth conditions

Strains and plasmids used in this study are listed in Supplementary Tables 4 and 5. *E. coli* strains were grown on Luria–Bertani agar (VWR) or in Luria–Bertani broth (VWR) with aeration at 37 °C. *S. aureus* strains were grown on tryptic soy agar (VWR), in tryptic soy broth (TSB, Difco) or in M9 minimal medium (KH_2_PO_4_ 3.4 g/l, VWR; K_2_HPO_4_ 2.9 g/l, VWR; di-ammonium citrate 0.7 g/l, Sigma-Aldrich; sodium acetate 0.26 g/l, Merck; glucose 1% (w/v), Merck; MgSO_4_ 0.7 mg/l, Sigma-Aldrich; CaCl_2_ 7 mg/l, Sigma-Aldrich; casamino acids 1% (w/v), Difco; MEM amino acids 1x, Thermo Fisher Scientific; MEM vitamins 1x, Thermo Fisher Scientific) at 200 rpm with aeration at 37°C, 30°C or 25°C. When necessary, culture media were supplemented with antibiotics (100 μg/ml ampicillin, Sigma-Aldrich; 10 μg/ml erythromycin, Apollo Scientific; 10 μg/ml chloramphenicol, Sigma-Aldrich). Unless otherwise specified, 5-bromo-4-chloro-3-indolyl β-d-galactopyranoside (X-Gal, Apollo Scientific) was used at 100 μg/ml, isopropyl β-d-1-thiogalactopyranoside (IPTG, Apollo Scientific) was used at 0.5 mM and anhydrotetracycline (Atc, Sigma-Aldrich) was used at 2 ng/ml.

### Construction of *S. aureus* strains

Oligonucleotides used in this study are listed in Supplementary Table 6. Cloning of FtsZ(T111A) mutants and tagged protein fusions in *S. aureus* was done using the following general strategy: plasmids were propagated in *E. coli* strains DC10B and purified from overnight cultures supplemented with the relevant antibiotics. Plasmids were then introduced into electrocompetent *S. aureus* RN4220 cells as previously described ^39^ and transduced to JE2 or COL using phage 80α ^40^. Antibiotic marker-free allelic replacements in the *S. aureus* chromosome were performed using plasmids that allow for double homologous recombination events at selected genome sites. Constructs were confirmed by PCR and by sequencing.

FtsZ(T111A) mutant strains were constructed using the pIMAY-Z vector ^41^. An *ftsZ* allele encoding the GTPase point mutation T111A (by substitution of ACT for GCA at bases 331-333) was generated by amplifying a 989-bp upstream and a 898-bp downstream region of codon 111 from the JE2 genome using the primers 6703/6726 and 6727/6700, respectively. The two fragments were joined by overlap PCR using the primers 6703/6700, digested with SmaI/SalI and cloned into SmaI/XhoI-digested pIMAY-Z. The resulting plasmid pIMAY-Z-ftsZ(T111A) was electroporated into RN4220 and transduced into JE2, JE2 EzrA-sGFP and COL EzrA-sGFP. Integration-excision by a double homologous recombination event was performed at 30°C, the *ftsZ* genomic region was sequenced to confirm the presence of the T111A mutation, and the chromosomal DNA of the JE2 EzrA-sGFP FtsZ(T111A) strain was sent for whole-genome sequencing.

Strains producing FtsW-HT, MurJ-HT, PBP4-HT, or GpsB-HT fusions from their respective native genomic locus were constructed by allelic replacement strategies using the pMAD vector ^42^. In brief, DNA fragments with approximately 800 bp spanning 3’-ends (excluding stop codons) of the *ftsW*, *murJ*, *pbp4* and *gpsB* genes from JE2 were amplified using the primers 7373/7142, 7376/6785, 7370/6783, and 7766/7597, respectively. A codon-optimized *halo-tag* sequence was synthesized (Integrated DNA Technologies, IDT; Supplementary Information) and amplified with the primer pairs 7031/7369 (for *ftsW-ht*), 6715/7369 (for *murJ-ht* and *pbp4-ht*) or 6715/7769 (for *gpsB-ht*). Corresponding DNA fragments were joined either by overlap PCR using the primers 7373/7369 (to generate *ftsW-ht*) or by SalI digestion followed by ligation (to generate *murJ-ht, pbp4-ht* and *gpsB-ht*). The downstream region of the *ftsW*, *murJ*, *pbp4* and *gpsB* genes from JE2 were amplified using the primers 7374/7375, 7377/7378, 7371/7372 and 7770/7771, respectively. Corresponding DNA fragments were joined either by KpnI digestion followed by ligation (for *ftsW-ht*, *murJ-ht*, and *pbp4-ht*) or by overlap PCR using the primers 7766/7771 (for *gpsB-ht*). The full constructs were then digested with EcoRI/BamHI (for *ftsW-ht, murJ-ht*, and *pbp4-ht*) or SmaI/BamHI (for *gpsB-ht*) and cloned into equally digested pMAD. Integration and excision of the pMAD derivatives in JE2 EzrA-sGFP and COL EzrA-sGFP derivative strains by a double homologous recombination event that led to allelic exchange was performed as previously described ^42^.

Strains ectopically producing HT-DivIB, RodA-HT, FtsW-HT, FtsZ-HT, EzrA-HT and iST-PBP1 were constructed using pBCB13 and pBCB43 vectors ^43, 44^, which are derivatives of pMAD that allow gene expression from the ectopic *spa* locus under the control of (IPTG-inducible) *spac* promoter and (Atc-inducible) *xyl-tetO* promoter, respectively. Briefly, a codon-optimized *halo-tag* sequence (Supplementary Information) was amplified with the primer pairs 6713/6714 and 6715/6716, digested with SmaI and cloned into equally digested pBCB13, to generate plasmids pBCB13-Nht and pBCB13-htC, respectively. The in-frame insertion of a coding sequence into the multiple cloning site of pBCB13-Nht or pBCB13-htC results in a translational fusion containing the HT sequence either at the gene product’s N- or C-terminus, respectively. The *divIB* full-length and truncated coding sequences (each excluding the start codon) were amplified from JE2 using primers 7590/7591 and 7590/9171, digested with XhoI/EagI and ligated into SalI/EagI-digested pBCB13-Nht, to generate pBCB13-htdivIB and pBCB13-htdivIB(Δγ), respectively. The *rodA* full-length coding sequence (excluding the stop codon) was amplified from JE2 using the primers 7594/7595, digested with EagI/SalI and ligated into equally digested pBCB13-htC, to generate pBCB13-rodAht. For the construction of pBCB43 derivatives, the *ftsW, ftsZ* and *ezrA* full-length coding sequences (excluding the stop codons) were amplified from JE2 using the primers 9247/7142, 7191/7267 and 7139/7140, respectively. Active-site mutant derivatives of *ftsW* were amplified from plasmids pCNX-ftsW(W121A)sgfp and pCNX-ftsW(D287A)sgfp ^18^ using the same *ftsW* primers. Note that primer 9247 contains a *tetO* sequence for an improved TetR repression. A codon-optimized *halo-tag* sequence was amplified with the primer pair 7031/7034 and joined with the *ftsW, ftsZ* and *ezrA* coding sequences by overlap PCR using the primer pairs 9247/7034, 7191/7034 and 7139/7034, respectively. The full constructs were then SmaI/EagI-digested and cloned into equally digested pBCB43. To generate pBCB43-istpbp1, the *pbp1* full-length coding sequence (excluding the start codon) was amplified from JE2 using the primer pair 3810/9647 and the *snap-tag* coding sequence amplified from pSNAP-tag (T7)-2 (New England Biolabs, NEB; Supplementary Information) using the primer pair 9631/9632. Note that primer 9631 adds an *i-tag* sequence (^45^; Supplementary Information) for elevated expression levels in *S. aureus*. The two DNA fragments were joined by overlap PCR using the primer pair 9631/9647, digested with SmaI/XbaI and ligated into SmaI/NheI-digested pBCB43. Integration and excision of the pBCB13 and pBCB43 derivatives in JE2 EzrA-sGFP and COL EzrA-sGFP derivative strains by a double homologous recombination event that led to gene replacement at the *spa* locus was performed as previously described ^43^.

### Growth curves of *S. aureus* strains

To assess growth of JE2 EzrA-sGFP derivative strains encoding ST and HT protein fusions, overnight cultures in TSB were back-diluted 1:1,000 into fresh media. A 200 µl sample of each culture was added to a well in a 96-well plate. Plates were incubated shaking at 37°C and the OD_600_ was recorded every 15 min for eight hours in a 96-well plate reader (Biotek Synergy Neo2). Cells producing HT-DivIB, EzrA-HT and iST-PBP1 were grown in presence of 0.5 mM IPTG, 0.5 ng/ml Atc and 2 ng/ml Atc, respectively.

To determine the cell growth rates of JE2 EzrA-sGFP, COL EzrA-sGFP and ColPBP1TP EzrA-sGFP derivative strains in fast and slow growth conditions, overnight cultures in TSB at 37°C, 30°C or 25°C were back-diluted to an OD_600_ of 0.01 in 50 ml TSB or M9 minimal medium, in 250-ml Erlenmeyer flasks. Cells were grown with shaking at 200 rpm at 37°C, 30°C or 25°C in triplicate. 1-ml samples were taken every 20 min, 30 min or 40 min to record the OD_600_ using a spectrophotometer (Biochrom Ultrospec 2100 Pro). Growth rates were calculated for each strain during its exponential growth phase.

### Protein in-gel fluorescence detection and Western blot analysis

To assess the integrity of ST and HT protein fusions, JE2 EzrA-sGFP derivative strains were grown to mid-exponential phase (OD_600_ of 0.6-0-8) in 50 ml TSB at 37°C. Cells producing ectopic HT-DivIB, EzrA-HT and iST-PBP1 were grown in presence of 0.5 mM IPTG, 0.5 ng/ml Atc and 2 ng/ml Atc, respectively. Cells producing FtsW-HT wild-type and mutant derivatives from the ectopic *spa* locus were grown in presence of 0.2 ng/ml Atc. Cells were harvested by centrifugation, resuspended in 0.3 ml fresh TSB and labelled with 500 nM of either JF549-HTL (red-fluorescent Janelia Fluor 549 Halo-tag ligand) or JF549-cpSTL (red-fluorescent Janelia Fluor 549 cell-permeable Snap-tag ligand) for 20 min at 37°C. Cells were cooled on ice, washed one time with 1 ml phosphate-buffered saline (PBS) and resuspended in 0.3 ml PBS supplemented with complete mini protease Inhibitor Cocktail (Roche). Cell suspensions were transferred to lysis tubes containing glass beads and subjected to mechanical disruption in a homogenizer SpeedMill Plus (Bioanalytik Jena) programmed to six 1-min cycles. Glass beads and cell debris were removed in two steps of centrifugation each for 1 min at 3,400 g. 25 µl of non-boiled protein sample were loaded on 12% Mini-Protean TGX pre-cast gels (Bio-Rad) and the proteins separated by SDS-PAGE. Gels were imaged in a FujiFilm FLA-5100 imaging system. EzrA-sGFP fluorescence was detected using 473-nm laser/Cy3 filter and JF549 fluorescence was detected using 532-nm laser/LPB filter settings.

To detect FtsZ wild-type and T111A mutant variants, JE2 EzrA-sGFP FtsW-HT and COL EzrA-sGFP FtsW-HT derivative strains were grown to mid-exponential phase (OD_600_ of 0.6-0-8) in 50 ml TSB at 30°C. Cells were cooled on ice, harvested by centrifugation and resuspended in 0.3 ml PBS supplemented with complete mini protease Inhibitor Cocktail (Roche). Whole cell protein extracts were obtained as described above and protein concentrations determined using Bradford reagent (Thermo Scientific). 2.5 μg of total protein were loaded on 12% Mini-Protean TGX pre-cast gels (Bio-Rad). Separated proteins were then transferred to a nitrocellulose membrane using a Trans-Blot Turbo RTA Mini 0.2 μm Nitrocellulose Transfer Kit and Trans-Blot Turbo system (Bio-Rad). The membrane was cut to separate the regions above and below approximately 70 kDa. The top part of the membrane was incubated with Sypro-Ruby stain (Invitrogen) following the manufacturer’s instructions to label high-molecular-weight proteins. The bottom part of the membrane containing FtsZ was blocked with 5% milk, followed by consecutive incubations with an anti-FtsZ antibody (1:2,000 dilution) for 16 h at 4°C and with a secondary fluorescent antibody (Alexa-488 anti-sheep diluted 1:50,000; Thermo Fisher) for 1 h at room temperature. Alexa-488 and Sypro-Ruby fluorescence detection was performed in an iBright Imaging System (Invitrogen).

### Transmission electron microscopy

Transmission electron microscopy (TEM) was essentially performed as previously described ^46^. Briefly, cells of strains JE2 and JE2 FtsZ(T111A) were grown in TSB rich medium at 37°C to mid-exponential phase (OD_600_ of 0.6-0.8) and harvested by centrifugation. Cell pellets were resuspended and fixed in wash buffer (0.1 M PIPES, pH 7.2) containing 2.5% glutaraldehyde (Carl Roth) and incubated on ice for one hour, followed by three wash steps with wash buffer. Cells were then post-fixed in wash buffer containing 1% osmium tetroxide (Acros Organics) for one hour at 4°C and washed five times with MilliQ H_2_O. Cells were embedded in 2% low melting point agarose, stained with 0.5% uranyl acetate (Analar) in H_2_O over night at 4°C, and washed twice with MilliQ H_2_O. Dehydration was performed at 4°C by increasing the ethanol concentration in samples gradually from 30% to 50%, 70%, 80%, 90% and 100% for 10 min in each step. The final step was repeated one time and ethanol exchanged with ice-cold acetone in two steps for 10 min and 20 min at room temperature. Infiltration and embedding were performed using Spurŕs resin (Science Services) and samples were polymerized for 24 to 48 hours at 60°C. Ultrathin sections (70 nm) were generated with an EM UC7 ultramicrotome (Leica), mounted on copper palladium slot grids coated with 1% formvar (Agar Scientific) in chloroform and post-stained with uranyl acetate and Reynold’s lead citrate for 5 min each. TEM imaging was performed at 120 kV using a FEI Tecnai G2 Spirit BioTWIN microscope equipped with an Olympus-SIS Veleta CCD Camera.

### Fluorescence microscopy

To study the localization of ST and HT protein fusions, JE2 EzrA-sGFP and COL EzrA-sGFP derivative strains were grown overnight in TSB and diluted 1:200 in fresh TSB followed by incubation with shaking at 37°C. For cells producing HT-DivIB, EzrA-HT and iST-PBP1 the medium was supplemented with 0.5 mM IPTG, 0.5 ng/ml Atc and 2 ng/ml Atc, respectively. After cells reached mid-exponential growth phase (OD_600_ of 0.6-0.8), 500 nM of either JF549-HTL or JF549-cpSTL were added to 1 ml of culture followed by incubation with shaking for 20 min at 37°C. Where indicated, cells were simultaneously treated with 2 µg/ml vancomycin (Sigma-Aldrich) for 20 min at 37°C. Cells were pelleted by centrifugation for 1 min at 9,300 g, washed one time, resuspended in PBS, and spotted on a microscope slide covered with a thin layer of 1.5% TopVision agarose (Thermo Fisher) in PBS. Images were acquired with a Zeiss Axio Observer microscope equipped with a Plan-Apochromat 100x/1.4 oil Ph3 objective, a Retiga R1 CCD camera (QImaging), a white-light source HXP 120 V (Zeiss) and the software ZEN blue (Zeiss). For image acquisition, the filters (Semrock) Brightline GFP-3035B (GFP) and Brightline TXRED-4040B (JF549) were used.

To analyse the cell size and cell cycle of *S. aureus*, overnight cultures in TSB of COL derivative strains were inoculated in quadruplicates from independent single colonies and incubated shaking at 30°C or 37°C. Cultures were diluted 1:200 into fresh TSB followed by incubation with shaking at the same temperatures. 1 ml of mid-exponential growth phase cells (OD_600_ of 0.6-0.8) was mixed with 5 µg/ml Nile red (Invitrogen) and 1 µg/ml Hoechst 33342 (Invitrogen) and incubated shaking for 5 min. Cells were washed one time with PBS and spotted on a pad of 1.5% TopVision agarose (Thermo Fisher) in PBS, and mounted in a Gene Frame (Thermo Fisher) on a microscope slide. Cells were then imaged by Structured Illumination Microscopy (SIM) using an Elyra PS.1 microscope (Zeiss) equipped with a Plan-Apochromat 63×/1.4 oil differential interference contrast M27 objective. SIM images were acquired using five grid rotations with 34 µm grating period for the 561 nm laser (100 mW, at 50% maximal power) and 23 µm grating period for the 405 nm laser (100 mW, at 100% maximal power). Images were captured using a PCO Edge 5.5 camera and reconstructed using ZEN software (black edition, 2012, version 8.1.0.484). Following SIM image reconstruction, cell size was measured, and cells classified according to their cell cycle phase (phase 1: prior to initiation of membrane constriction at mid cell; phase 2: ongoing membrane constriction for septum synthesis; phase 3: closed septum), using the software eHooke ^47^.

To evaluate localization of PG synthesis activity, FtsZ(T111A) derivative and wild-type strains of JE2 EzrA-sGFP FtsW-HT and COL EzrA-sGFP FtsW-HT were grown to mid-exponential growth phase (OD_600_ of 0.6-0.8) in TSB rich medium at 30°C and then dually labelled with 500 nM JF549-HTL for 20 min and with 25 µM fluorescent D-amino acid HADA for 10 min. Cells were washed one time with PBS and spotted on a pad of 1.5% TopVision agarose (Thermo Fisher) in PBS, and mounted in a Gene Frame (Thermo Fisher) on a microscope slide. Imaging was performed in a DeltaVision OMX SR microscope equipped with an Olympus 60X PlanApo N/1.42 oil differential interference contrast objective and two PCO Edge 5.5 sCMOS cameras (one for DIC, HADA and GFP; one for JF549 and JFX650). The software AcquireSRsoftWoRx (GE) was programmed to acquire Z-stacks of five images with a step size of 300 nm using a 488-nm laser (100 mW, at 20% maximal power), a 568-nm laser (100 mW, at 20% maximal power) and a 405-nm laser (100 mW, at 30% maximal power), each with an exposure time of 100 ms. The software SoftWorRx (v7.2.1) was used for maximum intensity projection (MIP) of five images from each Z-stack, fluorescence channel alignment and image deconvolution.

To determine EzrA-sGFP ring constriction rates, JE2 EzrA-sGFP and COL EzrA-sGFP derivative strains were grown in duplicate overnight in TSB and diluted 1:200 in fresh TSB followed by incubation with shaking at 37°C or 30°C. Exponentially growing cells (OD_600_ of 0.6-0.8) were harvested by centrifugation for 1 min at 9,300 g, resuspended in 30 µl fresh TSB, and spotted on a microscope slide covered with a thin layer of 1.5% TopVision agarose (Thermo Fisher) in TSB:PBS (1:1). Where indicated, 8 µg/ml DMPI or 5 µg/ml PC190723 were added to 1 ml of culture for 2 min at 37°C prior to cell harvest and cells were maintained in presence of antibiotic during washing and imaging. The time between the cells contacting the pad and the start of image acquisition was five minutes. Images were acquired in a DeltaVision OMX SR microscope (GE) in SIM mode. SIM images (three phase shifts and three grid rotations) were acquired every three minutes for 75 min (90 min for imaging at 30°C, 60 min for JE2 background) using a 488-nm laser (100 mW, at 10% maximal power, 5% for JE2 background) with an exposure time of 25 ms (50 ms for JE2 background). Images were reconstructed using SoftWoRx and aligned using NanoJ-Core drift correction ^48^. 1-pixel straight lines were drawn on constricting EzrA-sGFP rings to generate space-time kymographs in Fiji ^49^. Only cells visibly constricting over a minimum of five consecutive time frames and that completed septum closure during a recorded time series were used for analysis, except for DMPI samples where septum closure events were seldom.

To analyse EzrA-sGFP movement dynamics, JE2 EzrA-sGFP and COL EzrA-sGFP derivative strains were grown in duplicate overnight in TSB and diluted 1:200 in fresh TSB followed by incubation with shaking at 37°C or 30°C. Exponentially growing cells (OD_600_ of 0.6-0.8) were harvested by centrifugation for 1 min at 9,300 g, resuspended in 30 µl fresh TSB, and spotted on a pad of 1.5% molecular biology grade agarose (Bio-Rad) in M9 minimal medium mounted in a Gene Frame (Thermo Fisher) on a microscope slide. Where indicated, cells were treated with DMPI and PC190723 as described above. The time between the cells contacting the pad and the start of image acquisition was five minutes and image acquisition was done during a maximum period of 30 min. Imaging was performed in a DeltaVision OMX SR microscope equipped with a hardware-based focus stability (HW UltimateFocus) and an environmental control module set to 37°C or 30°C. Z-stacks of three images with a step size of 500 nm were acquired every three seconds for three minutes using a 488-nm laser (100 mW, at 10% maximal power) with an exposure time of 50 ms. MIP of three images from each Z-stack and subsequent image deconvolution was performed for each time frame in SoftWoRx. All 61 time frames were aligned using NanoJ-Core drift correction ^48^ and then used to perform MIP for the drawing of 1-pixel freehand lines over EzrA-sGFP in late pre-divisional cells, in which nascent Z-rings appear sparse and D-shaped. Space-time kymographs were then generated by extracting fluorescence intensities along drawn lines from individual time frames using the software Fiji ^49^. FtsZ treadmilling speed was calculated in nm/s by measuring the slopes of straight lines drawn on diagonals spanning the entire width in kymographs and corresponding to circumferentially moving EzrA-sGFP.

To perform single-molecule imaging, JE2 EzrA-sGFP and COL EzrA-sGFP derivative strains producing ST or HT protein fusions were grown in triplicate, overnight, in TSB and diluted 1:200 in fresh TSB or M9 minimal medium. For cells producing HT-DivIB, EzrA-HT and iST-PBP1 the medium was supplemented with 0.5 mM IPTG, 0.5 ng/ml Atc and 2 ng/ml Atc, respectively. Cells were grown with shaking at 37°C, 30°C or 25°C until mid-exponential growth phase (OD_600_ of 0.4-0.8). 1 ml of cell culture was then mixed either with 5 nM JFX650-STL (far-red-fluorescent Janelia Fluor X 650 Snap-tag ligand) or with JF549-HTL at concentrations ranging from 10 to 250 pM (Supplementary Tables 2 & 3), and incubated with shaking for 20 min at 37°C, 30°C or 25°C. Cells were harvested by centrifugation for 1 min at 9,300 g, resuspended in 30 µl fresh M9 minimal medium, spotted on a pad of 1.5% molecular biology grade agarose (Bio-Rad) in M9 minimal medium, mounted in a Gene Frame (Thermo Fisher) on a microscope slide, and covered with a glass coverslip pre-washed with changes of ethanol, acetone, 0.1 M KOH and MilliQ H_2_O. Where indicated, cells were treated with DMPI or PC190723 as described above, or with 10 µg/ml imipenem (Apollo Scientific) or 2 µg/ml vancomycin (Sigma-Aldrich), for 20 min at 37°C prior to harvest, and cells maintained in presence of antibiotic during washing and imaging. The time between the cells contacting the pad and the start of image acquisition was five minutes and image acquisition was done during a maximum period of 30 min. Imaging was performed in a DeltaVision OMX SR microscope equipped with a hardware-based focus stability (HW UltimateFocus) and an environmental control module set to 37°C, 30°C or 25°C. Z-stacks of three epifluorescence images with a step size of 500 nm were acquired every three seconds (or every second for a frame rate of 1 Hz) for three minutes (or for two minutes) using a 568-nm laser (100 mW, at 10% maximal power; for JF549-labelled FtsW-HT and HT-DivIB) or a 640-nm laser (100 mW, at 30% maximal power; for JFX650-labelled iST-PBP1), each with an exposure time of 800 ms (or 300 ms). Additional Z-stacks were acquired in the first and the last time frames of every time series to record EzrA-sGFP fluorescence using a 488-nm laser (100 mW, at 10% maximal power) with an exposure time of 100 ms and to image cells in brightfield. MIP of three images from each Z-stack acquired in both fluorescence channels and fluorescence channel alignment was performed for each time frame using SoftWoRx. 1-pixel freehand lines were drawn on the EzrA-sGFP signal in the last time frame and space-time kymographs were generated using Fiji ^49^ by extracting fluorescence intensities from recorded single molecules in all 61 time frames.

### Single-molecule tracking data analysis

Spots corresponding to fluorescence signal from single molecules of HT- and ST-labelled protein fusions were detected in TrackMate (v.7.2.0) ^50^ using the Laplacian of Gaussian (LoG) filter with sub-pixel localization, a blob diameter of 400 nm and a quality threshold of 3. Tracks were generated by linking spots detected in two consecutive time frames (3-s or 1-s interval) using the simple Linear Assignment Problem (LAP) tracker with a maximum linkage distance of 125 nm and no frame gaps allowed. Obtained tracks were filtered for a minimal number of spots of 30 for 0.33 Hz image acquisitions (equivalent to a duration of ≥87 s) or of 60 for 1 Hz image acquisitions (equivalent to a duration of ≥59 s). To calculate the percentage of cells containing a track, cells imaged in brightfield were counted using Otsu thresholding in Fiji ^49^. All further analysis was done by exporting the sub-pixel coordinates for each spot in a track from TrackMate to be used in an in-house developed Python tool (https://github.com/Bacterial CellBiologyLab/AureusSpeedTracker).

To calculate single-molecule motion in all three spatial dimensions of each detected molecule, an ellipse was manually drawn on the EzrA-sGFP signal for all cells with a directionally moving FtsW, PBP1, or DivIB molecule. The angle of the division ring relative to the image acquisition plane (Θ) was determined for all drawn ellipses by calculating the arccosine of the ratio between the minor axis, m, and major axis, M (equation 1). Each track was then approximated to a track overlaying the division ring, by computing, for each track point, the closest point on the ellipse and projecting it onto a 3D ring (Fig. 3a; Supplementary Movie 6).

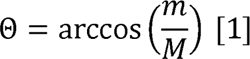

For each trajectory, *(x(t),y(t))* of length *L*, the Mean Squared Displacement (MSD) is calculated as a time average of the MSDs in each dimension ^51^:

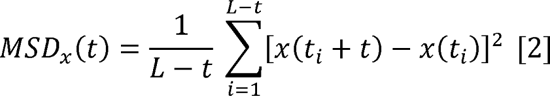

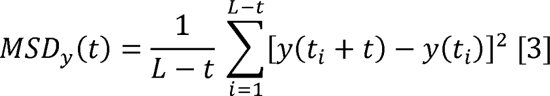

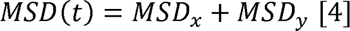

It is known that for pure Brownian motion the MSD is linear with time whilst for anomalous diffusion it follows a power law scaling. Taking o as a constant that depends on the diffusion coefficient and other constraints, and α is the anomalous diffusion exponent ^52^:

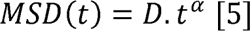

To calculate α the following linear regression was performed on the first 20 MSD points:

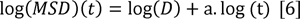

To determine single molecule velocities, the approximated tracks were unwrapped and sectioned by testing a maximum of three possible breakpoints (maximum of four sections). Possible sections and breakpoints were tested by performing linear regressions between every putative breakpoint, ensuring that all sections have a minimum length of four spots. The best possible sections, using 0 to 3 breakpoints, were selected by calculating the mean square error (MSE, equation 7) and choosing the combination of sections that has the minimum average MSE of all sections, weighted by section length. Velocities were calculated as the slope of the linear regression of each section. Unless mentioned otherwise, velocity refers to average velocity of all sections in a track.

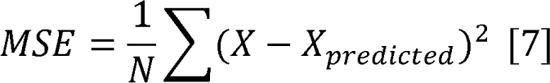

### *S. aureus* core divisome structure prediction and visualization

Predictions were generated for full-length *S. aureus* protein sequences using AlphaFold-Multimer ^53^ and AlphaFold2 ^54^ as implemented in ColabFold ^55^. All models for both methods produced similar predictions for the interface with PASTA domains (Cα RMSD 0.12–0.55 Å between PASTA domains in the top-ranked AlphaFold-Multimer model and the nine other models after alignment to DivIB residues 263–398). The top-ranked AlphaFold-multimer model is shown in Supplementary Fig. 12.

## Acknowledgements

We thank members of the Pinho lab, P. Pereira (ITQB-NOVA), S. Filipe (FCT-NOVA) and J. Xiao (Johns Hopkins University) for helpful discussions; L. Lavis (Janelia Research Campus, Ashburn) for the generous gift of JF549-HTL, JF549-cpSTL and JFX650-STL; E. Harry (University of Technology, Sydney) for providing the anti-FtsZ antibody; T. Roemer (Merck) for providing DMPI; M. S. VanNieuwenhze (Indiana University) for providing HADA; and the Electron Microscopy Facility and Genomics Unit of Instituto Gulbenkian de Ciencia. This study was funded by the European Research Council (ERC) through grant ERC-2017-CoG-771709 (to M.G.P.), by the European Union’s Horizon 2020 research and innovation programme under the Marie Sklodowska-Curie grant agreement N° 839596 (to S.S.), by the European Molecular Biology Organization (EMBO) through award ALTF 673-2018 (to S.S.), by The Company of Biologists Ltd (Journal of Cell Science) under the travelling fellowship agreement JCSTF1911323 (to S.S.), and by the Fundação para a Ciência e a Tecnologia through contract 2022.03033.CEECIND (to S.S.). R.H.’s contributions were supported by the Gulbenkian Foundation, the ERC (grant agreement N° 101001332), and the EMBO Installation Grant (EMBO-2020-IG-4734). Figure 7 was created with Biorender.com.

## Author contributions

S.S. and M.G.P. designed the research; S.S. performed the experiments; A.D.B. developed software with input from B.M.S., G.R.S., M.J.H., E.C.G. and R.H.; S.S. constructed strains; S.S., A.D.B. and M.G.P. analysed the overall data; Z.H. performed protein structure predictions; S.S. and M.G.P. wrote the manuscript with input from all authors.

