## Supplemental Tables and Figures for "Processive movement of *Staphylococcus aureus* essential septal peptidoglycan synthases is independent of FtsZ treadmilling and drives cell constriction"

Schäper *et al.*

*Supplementary Tables 1-6*

*Supplementary Figures 1-12*

*Supplementary Movies 1-8*

**Table S1. Overview of treadmilling speeds, septum constriction rates, and single-molecule velocities determined in this study.** Mean and standard deviation (SD) of EzrA-sGFP treadmilling speeds, EzrA-sGFP ring constriction rates, and single-molecule velocities of FtsW-HT and HT-DivIB. T, growth temperature. n.d., not determined.

|  |  | Treadmilling speed (nm/s) | |  | Septum constriction rate (nm/min) | |  | FtsW-HT velocity (nm/s) | |  | HT-DivIB velocity (nm/s) | |
| --- | --- | --- | --- | --- | --- | --- | --- | --- | --- | --- | --- | --- |
| T  (°C) | Genetic background, antibiotic treatment | Mean±SD | *n* |  | Mean±SD | *n* |  | Mean±SD | *n* |  | Mean±SD | *n* |
| 30 | JE2 | 40.2±9.0 | 171 |  | 14.6±3.8 | 20 |  | 10.6±3.0 | 623 |  | 11.0±3.3 | 897 |
|  | JE2 FtsZ(T111A) | 0.4±0.5 | 15 |  | 15.1±5.6 | 20 |  | 10.7±3.7 | 536 |  | 11.2±3.7 | 336 |
|  | COL | 39.6±8.5 | 83 |  | 18.0±4.6 | 20 |  | 9.4±3.6 | 262 |  | 9.6±3.3 | 131 |
|  | COL FtsZ(T111A) | 0.3±0.5 | 21 |  | 18.0±3.2 | 20 |  | 9.3±3.3 | 480 |  | 10.2±3.7 | 189 |
| 37 | JE2 | 60.6±9.2 | 32 |  | n.d. | |  | 15.7±4.3 | 486 |  | 16.5±5.0 | 194 |
|  | JE2, PC190723 | 0.4±0.5 | 9 |  | n.d. | |  | 16.0±4.8 | 449 |  | 16.4±4.4 | 282 |
|  | COL | 59.7±8.7 | 55 |  | 21.4±4.2 | 20 |  | 15.5±4.5 | 209 |  | 16.7±4.9 | 164 |
|  | COL, PC190723 | 0.3±0.5 | 6 |  | 22.5±3.0 | 20 |  | 14.0±4.4 | 230 |  | 15.3±4.6 | 144 |
|  | ColPBP1TP | 61.8±11.7 | 51 |  | 9.3±1.9 | 20 |  | 8.8±4.8 | 321 |  | 9.2±3.0 | 191 |
|  | COL, DMPI | 61.5±10.0 | 37 |  | 1.7±1.2 | 20 |  | n.d. | |  | n.d. | |
|  | JE2, DMPI | n.d. | |  | n.d. | |  | 4.9±2.6 | 227 |  | 6.4±3.0 | 343 |

**Table S2. FtsW, PBP1 and DivIB exhibit directional movement along septal rings as opposed to other cell division and cell wall-related proteins.** Tracks statistics for various cell division and cell wall-related proteins imaged by single-molecule tracking microscopy. Indicated strains were grown in TSB at 37°C and cells producing iST-PBP1, EzrA-HT or HT-DivIB derivatives from the ectopic *spa* locus were grown in presence of 2 ng/ml Atc, 0.5 ng/ml Atc or 0.5 mM IPTG, respectively. Cell division and cell wall-related proteins fused to Halo-tag (HT) were labelled with the red fluorescent dye JF549-HTL at indicated concentrations. iST-PBP1 was labelled with the far-red fluorescent dye JFX650-STL. Spot detection and tracking was performed in TrackMate (max. linkage distance, 125 nm; no frame gaps allowed). Obtained tracks were filtered for ≥30 spots per track (equivalent to a duration of ≥87 s). Only tracks overlapping with an EzrA-sGFP ring and an α_MSD_ ≥1 were included in this analysis. The number of tracks was normalized to the number of imaged cells (min. 1194) for each sample. Θ indicates the angle between the imaging and cell division planes.

| Strain | Dye conc. (pM) | Number  of tracks | % of cells with track | Avg. α_MSD_ | Avg. track  duration (s) | Avg. Θ (deg) |
| --- | --- | --- | --- | --- | --- | --- |
| JE2 EzrA-sGFP | 10 | 0 | 0.00 | - | - | - |
| JE2 EzrA-sGFP FtsW-HT | 10 | 125 | 5.13 | 1.48 | 120 | 51 |
| JE2 EzrA-sGFP MurJ-HT | 10 | 7 | 0.29 | 1.15 | 121 | 49 |
| JE2 EzrA-sGFP PBP4-HT | 10 | 0 | 0.00 | - | - | - |
| JE2 EzrA-sGFP GpsB-HT | 10 | 0 | 0.00 | - | - | - |
| JE2 EzrA-sGFP spa-FtsZ-HT | 10 | 0 | 0.00 | - | - | - |
| JE2 EzrA-sGFP spa-EzrA-HT | 10 | 0 | 0.00 | - | - | - |
| JE2 EzrA-sGFP spa-HT-DivIB | 10 | 22 | 1.23 | 1.51 | 132 | 53 |
| JE2 EzrA-sGFP spa-HT-DivIB | 50 | 63 | 3.12 | 1.51 | 112 | 53 |
| JE2 EzrA-sGFP spa-HT-DivIB(Δγ) | 50 | 1 | 0.03 | 1.22 | 105 | 31 |
| JE2 EzrA-sGFP spa-RodA-HT | 10 | 1 | 0.04 | 1.27 | 102 | 72 |
| JE2 EzrA-sGFP spa-Pxyl-tetO | 5,000 | 0 | 0.00 | - | - | - |
| JE2 EzrA-sGFP spa-iST-PBP1 | 5,000 | 192 | 0.69 | 1.45 | 114 | 55 |

**Table S3. Tracks statistics for FtsW and DivIB single molecules.** Indicated strains were grown in various conditions for single-molecule tracking microscopy. Cells producing HT-DivIB or FtsW-HT derivatives from the ectopic *spa* locus were grown in presence of 0.5 mM IPTG or 0.2 ng/ml Atc, respectively. HT-DivIB and FtsW-HT derivatives were labelled with the red fluorescent dye JF549-HTL. Tracking was performed in TrackMate (max. linkage distance, 125 nm; no frame gaps allowed). All tracks on top of EzrA-sGFP rings were filtered for number of spots in track (≥30, equivalent to ≥87 s) and α_MSD_ (≥1). The number of tracks was normalized to the number of imaged cells (min. 3374) for each sample and tracks were obtained from at least three biological replicates. Θ indicates the angle between the imaging and cell division planes.

| Fusion | Figure | Genetic background,  growth temperature,  growth medium/  antibiotic treatment/  protein variant | Dye conc. (pM) | Number  of tracks | % of cells with track | Avg. α_MSD_ | Avg. track duration (s) | Avg. Θ (deg) |
| --- | --- | --- | --- | --- | --- | --- | --- | --- |
| FtsW-HT | 4a | JE2, 37°C, TSB | 10 | 694 | 9.26 | 1.53 | 126 | 49 |
|  |  | JE2, 37°C, M9 | 50 | 472 | 7.13 | 1.48 | 131 | 64 |
|  |  | JE2, 30°C, TSB | 10 | 685 | 9.91 | 1.43 | 141 | 56 |
|  |  | JE2, 25°C, TSB | 10 | 385 | 4.14 | 1.36 | 148 | 46 |
|  | 5a | JE2, 30°C | 10 | 774 | 10.17 | 1.49 | 137 | 49 |
|  |  | JE2 FtsZ(T111A), 30°C | 10 | 597 | 12.34 | 1.48 | 131 | 50 |
|  |  | COL, 30°C | 20 | 344 | 4.65 | 1.41 | 137 | 54 |
|  |  | COL FtsZ(T111A), 30°C | 20 | 530 | 6.29 | 1.42 | 134 | 50 |
|  | 5c | JE2, 37°C | 10 | 619 | 7.63 | 1.52 | 127 | 53 |
|  |  | JE2, 37°C, PC190723 | 10 | 560 | 6.41 | 1.51 | 126 | 53 |
|  |  | COL, 37°C | 20 | 230 | 3.01 | 1.48 | 130 | 53 |
|  |  | COL, 37°C, PC190723 | 20 | 251 | 3.39 | 1.45 | 128 | 47 |
|  | 6a | JE2, 37°C | 10 | 619 | 7.63 | 1.52 | 127 | 53 |
|  |  | JE2, 37°C, imipenem | 10 | 397 | 8.29 | 1.24 | 133 | 47 |
|  |  | JE2, 37°C, DMPI | 10 | 243 | 4.46 | 1.27 | 135 | 54 |
|  |  | JE2, 37°C, vancomycin | 10 | 5 | 0.10 | 1.19 | 105 | 24 |
|  | 6c | COL, 37°C | 20 | 230 | 3.01 | 1.48 | 130 | 53 |
|  |  | ColPBP1TP, 37°C | 20 | 290 | 3.92 | 1.25 | 131 | 45 |
| FtsW-HT(spa) | 6e | JE2, 37°C, FtsW | 200 | 250 | 3.76 | 1.52 | 121 | 57 |
|  |  | JE2, 37°C, FtsW(W121A) | 200 | 59 | 0.38 | 1.37 | 116 | 48 |
|  |  | JE2, 37°C, FtsW(D287A) | 200 | 121 | 0.90 | 1.32 | 111 | 53 |
| HT-DivIB | 4b | JE2, 37°C, TSB | 50 | 499 | 5.72 | 1.51 | 120 | 52 |
|  |  | JE2, 37°C, M9 | 250 | 92 | 1.52 | 1.50 | 129 | 62 |
|  |  | JE2, 30°C, TSB | 50 | 989 | 12.14 | 1.44 | 137 | 59 |
|  |  | JE2, 25°C, TSB | 50 | 209 | 2.59 | 1.39 | 152 | 47 |
|  | 5b | JE2, 30°C | 50 | 1,073 | 12.10 | 1.50 | 131 | 48 |
|  |  | JE2 FtsZ(T111A), 30°C | 50 | 386 | 8.20 | 1.52 | 127 | 51 |
|  |  | COL, 30°C | 100 | 189 | 3.11 | 1.48 | 135 | 61 |
|  |  | COL FtsZ(T111A), 30°C | 100 | 218 | 2.90 | 1.46 | 136 | 53 |
|  | 5d | JE2, 37°C | 50 | 250 | 5.89 | 1.50 | 120 | 56 |
|  |  | JE2, 37°C, PC190723 | 50 | 345 | 5.05 | 1.51 | 118 | 54 |
|  |  | COL, 37°C | 100 | 217 | 2.30 | 1.50 | 124 | 58 |
|  |  | COL, 37°C, PC190723 | 100 | 181 | 1.97 | 1.48 | 125 | 56 |
|  | 6b | JE2, 37°C | 50 | 250 | 5.89 | 1.50 | 120 | 56 |
|  |  | JE2, 37°C, imipenem | 50 | 386 | 7.77 | 1.32 | 133 | 45 |
|  |  | JE2, 37°C, DMPI | 50 | 500 | 7.85 | 1.39 | 133 | 58 |
|  |  | JE2, 37°C, vancomycin | 50 | 9 | 0.19 | 1.17 | 119 | 35 |
|  | 6d | COL, 37°C | 100 | 217 | 2.30 | 1.50 | 124 | 58 |
|  |  | ColPBP1TP, 37°C | 100 | 178 | 2.24 | 1.32 | 132 | 45 |

**Table S4. Bacterial strains used in this study.**

| Name | Description | Reference |
| --- | --- | --- |
| *Escherichia coli* |  |  |
| DC10B | Δ*dcm* in DH10B background; Dam methylation only; for cloning | ^1^ |
| *Staphylococcus aureus* |  |  |
| RN4220 | Restriction-negative derivative of NCTC8325-4 | ^2^ |
| USA300 JE2 | CA-MRSA | ^3^ |
| JE2 FtsZ(T111A) | USA300 JE2 *ftsZ::ftsZ_T111A_* | This study |
| JE2 EzrA-sGFP | JE2 *ezrA::ezrA-sgfp;* CA-MRSA | ^4^ |
| JE2 EzrA-sGFP FtsZ(T111A) | JE2 *ezrA::ezrA-sgfp ftsZ::ftsZ_T111A_* | This study |
| COL EzrA-sGFP | COL *ezrA::ezrA-sgfp;* HA-MRSA | ^4^ |
| COL EzrA-sGFP FtsZ(T111A) | COL *ezrA::ezrA-sgfp ftsZ::ftsZ_T111A_* | This study |
| ColPBP1TP | COL *pbp1::pbp1_S314A_* | ^5^ |
| ColPBP1TP EzrA-sGFP | COL *pbp1::pbp1_S314A_ ezrA::ezrA-sgfp* | This study |
| JE2 EzrA-sGFP FtsW-HT | JE2 *ezrA::ezrA-sgfp ftsW::ftsW-halo* | This study |
| JE2 EzrA-sGFP FtsW-HT FtsZ(T111A) | JE2 *ezrA::ezrA-sgfp ftsW::ftsW-halo ftsZ::ftsZ_T111A_* | This study |
| COL EzrA-sGFP FtsW-HT | COL *ezrA::ezrA-sgfp ftsW::ftsW-halo* | This study |
| COL EzrA-sGFP FtsW-HT FtsZ(T111A) | COL *ezrA::ezrA-sgfp ftsW::ftsW-halo ftsZ::ftsZ_T111A_* | This study |
| ColPBP1TP EzrA-sGFP FtsW-HT | COL *pbp1::pbp1_S314A_ ezrA::ezrA-sgfp ftsW::ftsW-halo* | This study |
| JE2 EzrA-sGFP spa-HT-DivIB | JE2 *ezrA::ezrA-sgfp* Δ*spa::*P*_spac_-halo-divIB* | This study |
| JE2 EzrA-sGFP spa-HT-DivIB FtsZ(T111A) | JE2 *ezrA::ezrA-sgfp* Δ*spa::*P*_spac_-halo-divIB ftsZ::ftsZ_T111A_* | This study |
| COL EzrA-sGFP spa-HT-DivIB | COL *ezrA::ezrA-sgfp* Δ*spa::*P*_spac_-halo-divIB* | This study |
| COL EzrA-sGFP spa-HT-DivIB FtsZ(T111A) | COL *ezrA::ezrA-sgfp* Δ*spa::*P*_spac_-halo-divIB ftsZ::ftsZ_T111A_* | This study |
| ColPBP1TP EzrA-sGFP spa-HT-DivIB | COL *pbp1::pbp1_S314A_ ezrA::ezrA-sgfp* Δ*spa::*P*_spac_-halo-divIB* | This study |
| JE2 EzrA-sGFP MurJ-HT | JE2 *ezrA::ezrA-sgfp murJ::murJ-halo* | This study |
| JE2 EzrA-sGFP PBP4-HT | JE2 *ezrA::ezrA-sgfp pbp4::pbp4-halo* | This study |
| JE2 EzrA-sGFP GpsB-HT | JE2 *ezrA::ezrA-sgfp gpsB::gpsB-halo* | This study |
| JE2 EzrA-sGFP spa-Pxyl-tetO | JE2 *ezrA::ezrA-sgfp* Δ*spa::*P*_xyl-tetO_* | This study |
| JE2 EzrA-sGFP spa-FtsZ-HT | JE2 *ezrA::ezrA-sgfp* Δ*spa::*P*_xyl-tetO_-ftsZ-halo* | This study |
| JE2 EzrA-sGFP spa-EzrA-HT | JE2 *ezrA::ezrA-sgfp* Δ*spa::*P*_xyl-tetO_-ezrA-halo* | This study |
| JE2 EzrA-sGFP spa-FtsW-HT | JE2 *ezrA::ezrA-sgfp* Δ*spa::*P*_xyl-tetO_-ftsW-halo* | This study |
| JE2 EzrA-sGFP spa-FtsW(W121A)-HT | JE2 *ezrA::ezrA-sgfp* Δ*spa::*P*_xyl-tetO_-ftsW_W121A_-halo* | This study |
| JE2 EzrA-sGFP spa-FtsW(D287A)-HT | JE2 *ezrA::ezrA-sgfp* Δ*spa::*P*_xyl-tetO_-ftsW_D287A_-halo* | This study |
| JE2 EzrA-sGFP spa-HT-DivIB(Δγ) | JE2 *ezrA::ezrA-sgfp* Δ*spa::*P*_spac_-halo-divIB_2-372_* | This study |
| JE2 EzrA-sGFP spa-iST-PBP1 | JE2 *ezrA::ezrA-sgfp* Δ*spa::*P*_xyl-tetO_-snap-pbp1* | This study |
| JE2 EzrA-sGFP spa-RodA-HT | JE2 *ezrA::ezrA-sgfp* Δ*spa::*P*_spac_-rodA-halo* | This study |

**Table S5. Plasmids used in this study.**

| Name | Description | Reference |
| --- | --- | --- |
| pSNAP-tag (T7)-2 | E. coli expression vector encoding the Snap-tag protein; Amp^r^ | NEB |
| pIMAY-Z | *E. coli-S. aureus* shuttle vector with a thermosensitive origin of replication for Gram positive bacteria; Cm^r^, *lacZ* | ^6^ |
| pIMAY-Z-ftsZ(T111A) | pIMAY-Z derivative containing *ftsZ_T111A_*; Cm^r^, *lacZ* | This study |
| pMAD | *E. coli-S. aureus* shuttle vector with a thermosensitive origin of replication for Gram positive bacteria; Amp^r^, Ery^r^, *lacZ* | ^7^ |
| pMAD-ezrAsgfp | pMAD derivative containing an *ezrA-sgfp* fusion and the downstream region of *ezrA;* Amp^r^, Ery^r^ | ^5^ |
| pMAD-ftsWht | pMAD derivative containing an *ftsW-halo* fusion and the downstream region of *ftsW;* Amp^r^, Ery^r^ | This study |
| pMAD-murJht | pMAD derivative containing a *murJ-halo* fusion and the downstream region of *murJ;* Amp^r^, Ery^r^ | This study |
| pMAD-pbp4ht | pMAD derivative containing a *pbp4-halo* fusion and the downstream region of *pbp4;* Amp^r^, Ery^r^ | This study |
| pMAD-gpsBht | pMAD derivative containing a *gpsB-halo* fusion and the downstream region of *gpsB;* Amp^r^, Ery^r^ | This study |
| pCNX-ftsW(W121A)sgfp | pCNX derivative containing an *ftsW_W121A_-sgfp* fusion; Amp^r^, Kan^r^ | ^5^ |
| pCNX-ftsW(D287A)sgfp | pCNX derivative containing an *ftsW_D287A_-sgfp* fusion; Amp^r^, Kan^r^ | ^5^ |
| pBCB43 | pMAD derivative with up- and downstream regions of the *spa* locus and *tetR-*P*_xyl-tetO_*; Amp^r^, Ery^r^, *lacZ* | ^8^ |
| pBCB43-ftsZht | pBCB43 derivative containing an *ftsZ-halo* fusion; Amp^r^, Ery^r^, *lacZ* | This study |
| pBCB43-ezrAht | pBCB43 derivative containing an *ezrA-halo* fusion; Amp^r^, Ery^r^, *lacZ* | This study |
| pBCB43-ftsWht | pBCB43 derivative containing an *ftsW-halo* fusion; Amp^r^, Ery^r^, *lacZ* | This study |
| pBCB43-ftsW(W121A)ht | pBCB43 derivative containing an *ftsW_W121A_-halo* fusion; Amp^r^, Ery^r^, *lacZ* | This study |
| pBCB43-ftsW(D287A)ht | pBCB43 derivative containing an *ftsW_D287A_-halo* fusion; Amp^r^, Ery^r^, *lacZ* | This study |
| pBCB43-istpbp1 | pBCB43 derivative containing a *isnap-pbp1* fusion; Amp^r^, Ery^r^, *lacZ* | This study |
| pBCB13 | pMAD derivative with up- and downstream regions of the *spa* locus and *lacI-*P*_spac_*; Amp^r^, Ery^r^, *lacZ* | ^9^ |
| pBCB13-Nht | pBCB13 derivative containing *halo-tag* for N-terminal protein fusions; Amp^r^, Ery^r^, *lacZ* | This study |
| pBCB13-htdivIB | pBCB13-htN derivative containing a *halo-divIB* fusion; Amp^r^, Ery^r^, *lacZ* | This study |
| pBCB13-htdivIB(Δγ) | pBCB13-htdivIB derivative encoding Halo-DivIB C-terminally truncated by 67 aa; Amp^r^, Ery^r^, *lacZ* | This study |
| pBCB13-htC | pBCB13 derivative containing *halo-tag* for C-terminal protein fusions; Amp^r^, Ery^r^, *lacZ* | This study |
| pBCB13-rodAht | pBCB13-htC derivative containing a *rodA-halo* fusion; Amp^r^, Ery^r^, *lacZ* | This study |

**Table S6. Oligonucleotides used in this study.**

| No. | Name | 5ʹ-3ʹ sequence |
| --- | --- | --- |
| 3810 | 10aa linker-PBP1 for | GGCGGTTCTGGCGGAGGTGGCTCTGCGAAGCAAAAAATTAAAATTAAA |
| 6700 | COLftsZ_stop+58_rev | ATATCCCGGGCATCAGATATGTTATCTGATGATTTGT |
| 6703 | COLftsZ_-659_fwd | ATATGTCGACGATTCTGCTTCAGATCAAGATATCTTC |
| 6713 | ftsZ-RBS_fwd | ATATCCCGGGGGCCAATAAAACTAGGAGGAAATTTAA |
| 6714 | 10aa-linker_rev | ATATCCCGGGAGTACTCGGCCGGTCGACAGAGCCACCTCCGCCAGAACCGCCTCCACC |
| 6715 | 5aa-linker-C-halo-tag_fwd | ATATCCCGGGAGTACTCGGCCGGTCGACTCCTGCGGCGCCTCCGCGGAGATTGGAACTGGTTT |
| 6716 | C-halo-tag_rev | ATATCCCGGGTTAACCACTGATTTCTAAAGTAGATAACCATC |
| 6726 | COLftsZ-T111A_rev | AACGACTGGTGCTGCACCTGCACCAGTTCCGCCACCCATA |
| 6727 | COLftsZ-T111A_fwd | GCAGGTGCAGCACCAGTCGTT |
| 6783 | COLpbp4_-stop_rev | ATATGTCGACTTTTCTTTTTCTAAATAAACGATTGA |
| 6785 | COLmurJ_-stop_rev | ATATGTCGACTCGTAAAAACCTAACTCTACGTCTT |
| 7031 | C-HT_15aa_fwd | TTGGAAGGATCAGGACAAGGACCAGGATCTGGTCAAGGTTCTGGTGCGGAGATTGGAACTGGTTTCCCGT |
| 7034 | C-halo-tag_rev2 | ATATCTCGAGCGGCCGTTAACCACTGATTTCTAAAGTAGATAACCATC |
| 7139 | COLezrA_st7_SmaI_fwd | ATATCCCGGGAAAAAATAAGGAGGAAAAAAAATGGTGTTATATATCATTTTGGCAAT |
| 7140 | COLezrA-15aa_-stop_rev | AGATCCTGGTCCTTGTCCTGATCCTTCCAATTGCTTAATAACTTCTTCTTCAATA |
| 7142 | COLftsW-15aa_-stop_rev | AGATCCTGGTCCTTGTCCTGATCCTTCCAAATTAAATTGTCTTCTTATATCAAC |
| 7191 | COLftsZ+1_st7_fwd | ATATCCCGGGAAAAAATAAGGAGGAAAAAAAATGTTAGAATTTGAACAAGGATTTAAT |
| 7267 | ftsZ-15aa_rev2 | AGATCCTGGTCCTTGTCCTGATCCTTCCAAACGTCTTGTTCTTCTTGAACGTCTT |
| 7369 | HT_Kpn_rev | ATATGGTACCTTAACCACTGATTTCTAAAGTAG |
| 7370 | pbp4_800int_Eco_fwd | ATATGAATTCTTCGTCAATCCAACGGGTGCTG |
| 7371 | pbp4_800down_Kpn_fwd | ATATGGTACCAACATACTAAAAACGGACAAGTTGC |
| 7372 | pbp4_800down_Bam_rev | ATATGGATCCACCCAGCAGTAACGCACACGACAAT |
| 7373 | ftsW_800int_Eco_fwd | ATATGAATTCATGAACTTACAGGCATCTGAGT |
| 7374 | ftsW_800down_Kpn_fwd | ATATGGTACCAAAAATACTAGCCAATATTTAG |
| 7375 | ftsW_800down_Bam_rev | ATATGGATCCACGACGCGCAAATTGTTCATT |
| 7376 | murJ_800int_Bam_fwd | ATATGGATCCTACCTTCACAGTTACAAGATATATT |
| 7377 | murJ_800down_Kpn_fwd | ATATGGTACCTTAAGACGTAGAGTTAGGTT |
| 7378 | murJ_800down_Eco_rev | ATATGAATTCTCACGACTATCTTTACGTGTAACAAGT |
| 7590 | divIB_+4_Xho_fwd | ATATCTCGAGGATGATAAAACGAAGAACGATCAACA |
| 7591 | divIB_EagI_rev | ATATCGGCCGTTAATTATTCTTACTTGATTGTTTGT |
| 7594 | rodA_Eag_st7_fwd | ATATCGGCCGAAAAAATAAGGAGGAAAAAAAATGAATTATTCATCTCGTCAACAGCCG |
| 7595 | rodA_-stop_Sal_rev | ATATGTCGACATTACTTTTTGGATGGTATAAATCGA |
| 7597 | gpsB_-stop_Sal_rev | ATATGTCGACTTTACCAAATACAGCTTTTTCTAAGTTT |
| 7766 | GA_gpsB-HT_1 | GCATGCCATGGTACCCGGGAGCTCGAATTCATGGTTAAAACAGTTTATGTAACAG |
| 7769 | GA_gpsB-HT_4 | TTGTATTTAGTAATTAACCACTGATTTCTAAAGTAG |
| 7770 | GA_gpsB-HT_5 | ATCAGTGGTTAATTACTAAATACAAAAGTTTAACTGTC |
| 7771 | GA_gpsB-HT_6 | TCCAGCCTCGCGTCGGGCGATATCGGATCCGTTCAACAATAGCTTTCTTAGTTATC |
| 9171 | divIB-372_EagI_rev | ATATCGGCCGTTATGATAATGATTGTGACATCTG |
| 9247 | ftsW_tetO-st3_fwd | TCTATCATTGATAGAGTCCCGGGAGCTCTCTATCATTGATAGAGTAAAAAAAAATAAGGAGGAAAATGAAGAATTTTAGAAGTATTTTACG |
| 9631 | iSNAP_st7_Sma_fwd | ATATCCCGGGAAAAAATAAGGAGGAAAAAAAATGGATAAAAAAGGTTTGGAAATTTTTTTGGCTTCTGACAAAGATTGCGAAATGAAACG |
| 9632 | SNAP-10aa_rev | AGAGCCACCTCCGCCAGAACCGCCTCCACC*TCCCAGACCCGGTTTACCCAG* |
| 9647 | 066-PBP1-stp-xbai | GGGCCCTCTAGATACGGCCGTTATTAGTCCGACTTATCCTTGTCAGTTTTAC |

Underlines sequences correspond to restriction sites used for cloning.

*halo-tag* sequence (IDT)

ATGGCGGAGATTGGAACTGGTTTCCCGTTCGACCCTCACTACGTGGAGGTCTTAGGAGAACGTATGCACTACGTGGACGTCGGTCCTCGTGATGGAACTCCAGTATTGTTCTTACATGGAAACCCTACTAGTTCATACGTGTGGCGAAACATCATACCTCATGTGGCACCGACACACCGTTGTATCGCTCCTGATTTAATCGGAATGGGCAAAAGTGACAAGCCGGACTTAGGTTATTTCTTTGACGATCACGTGCGATTTATGGATGCATTTATTGAGGCATTAGGATTAGAAGAAGTTGTCTTGGTAATACATGATTGGGGCTCTGCATTGGGCTTCCACTGGGCGAAGAGAAACCCTGAGCGAGTAAAGGGCATCGCGTTCATGGAGTTTATTCGTCCAATCCCAACATGGGATGAATGGCCGGAATTTGCTAGAGAAACGTTCCAAGCATTCCGTACTACTGACGTAGGCCGAAAGTTAATCATTGACCAGAACGTATTCATCGAGGGAACGTTACCTATGGGAGTGGTTAGACCATTGACAGAAGTCGAGATGGACCATTACCGAGAGCCGTTTTTGAACCCAGTGGACAGAGAGCCGTTATGGCGATTCCCTAACGAGTTACCTATTGCAGGTGAGCCGGCTAACATTGTCGCTTTGGTTGAAGAATATATGGACTGGTTGCATCAATCACCGGTCCCAAAATTATTGTTTTGGGGAACACCTGGTGTGTTGATCCCACCGGCTGAAGCTGCGCGTTTGGCGAAATCTTTGCCGAACTGTAAGGCGGTTGATATAGGTCCGGGATTAAACTTATTACAGGAGGACAATCCTGACTTAATAGGCAGTGAGATCGCTAGATGGTTATCTACTTTAGAAATCAGTGGT

*i-tag* sequence ^10^

ATGGATAAAAAAGGTTTGGAAATTTTTTTGGCTTCT

*snap-tag* sequence (NEB)

ATGGACAAAGATTGCGAAATGAAACGTACCACCCTGGATAGCCCGCTGGGCAAACTGGAACTGAGCGGCTGCGAACAGGGCCTGCATGAAATTAAACTGCTGGGTAAAGGCACCAGCGCGGCCGATGCGGTTGAAGTTCCGGCCCCGGCCGCCGTGCTGGGTGGTCCGGAACCGCTGATGCAGGCGACCGCGTGGCTGAACGCGTATTTTCATCAGCCGGAAGCGATTGAAGAATTTCCGGTTCCGGCGCTGCATCATCCGGTGTTTCAGCAGGAGAGCTTTACCCGTCAGGTGCTGTGGAAACTGCTGAAAGTGGTTAAATTTGGCGAAGTGATTAGCTATCAGCAGCTGGCGGCCCTGGCGGGTAATCCGGCGGCCACCGCCGCCGTTAAAACCGCGCTGAGCGGTAACCCGGTGCCGATTCTGATTCCGTGCCATCGTGTGGTTAGCTCTAGCGGTGCGGTTGGCGGTTATGAAGGTGGTCTGGCGGTGAAAGAGTGGCTGCTGGCCCATGAAGGTCATCGTCTGGGTAAACCGGGTCTGGGA

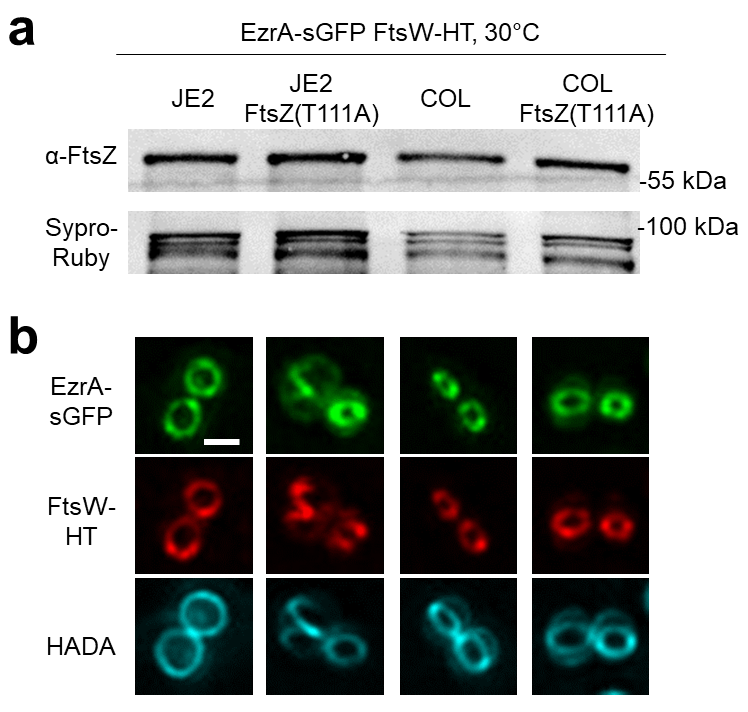

**Figure S1. *S. aureus* FtsZ GTPase mutants maintain native FtsZ levels and incorporation of septal peptidoglycan**. **a,** Western blot analysis of indicated strains grown in TSB rich medium at 30°C using anti-FtsZ antibody. Sypro-Ruby staining of immobilized proteins served as loading control. The theoretical molecular weight of FtsZ is 41.0 kDa. **b,** Representative epifluorescence micrographs of strains indicated in panel **a** labelled with JF549-HTL to visualize FtsW-HT and fluorescent D-amino acid HADA to visualize sites of nascent peptidoglycan synthesis. Scale bar, 0.5 µm.

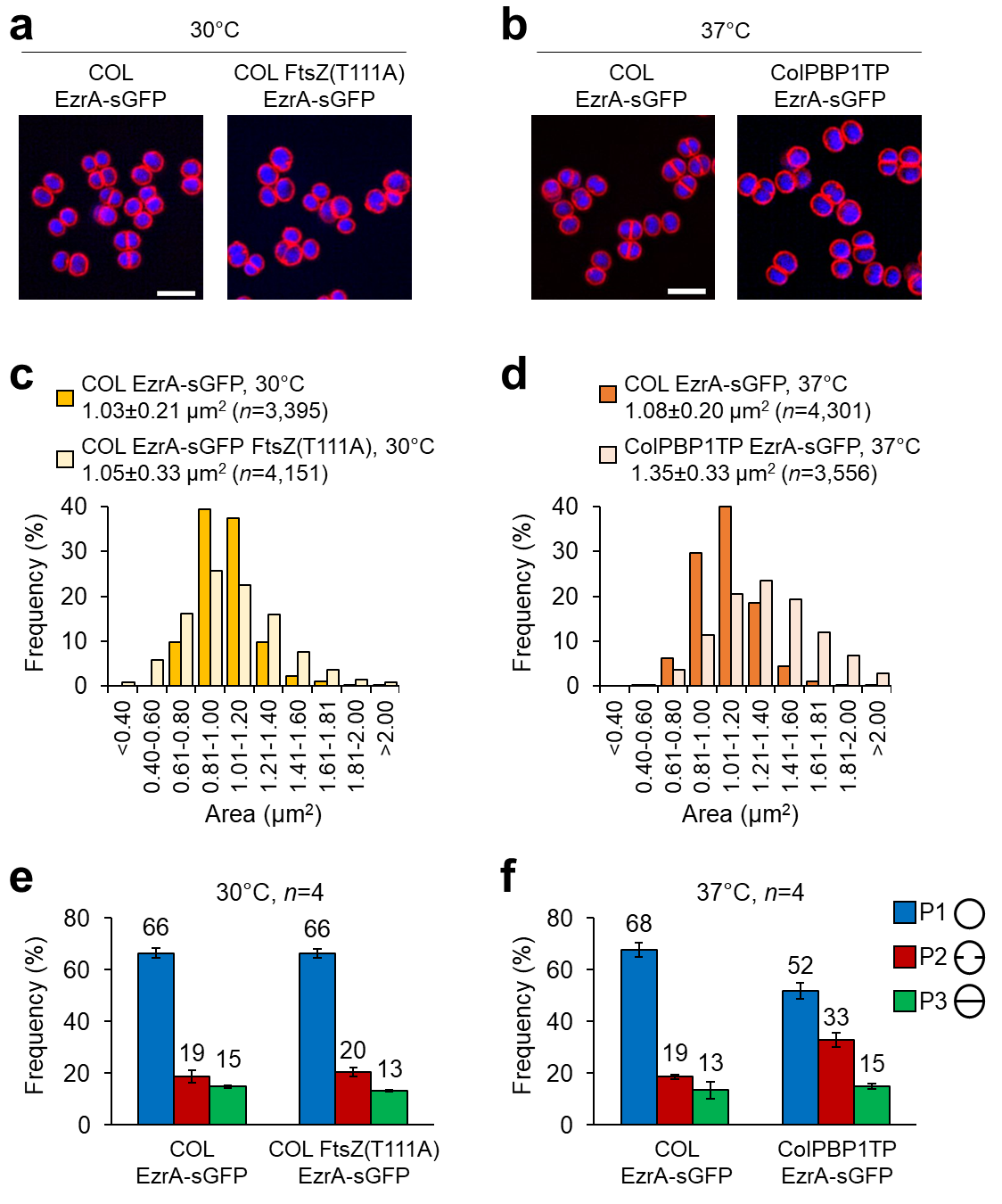

**Figure S2. *S. aureus* FtsZ GTPase and PBP1 TPase mutants are differentially affected in cell size and cell cycle progression. a,b,** Representative structured illumination micrographs of indicated strains grown to mid-exponential phase in TSB rich medium at indicated temperatures and labelled with the fluorescent dyes Nile red (membrane) and Hoechst 33342 (DNA). Scale bars, 2 µm. **c-f,** Determination of cell area (**c,d**) and classification of cells into three cell-cycle phases (**e,f**) for strains shown in panels **a** and **b**. Error bars represent the standard deviations for the mean from four biological replicates.

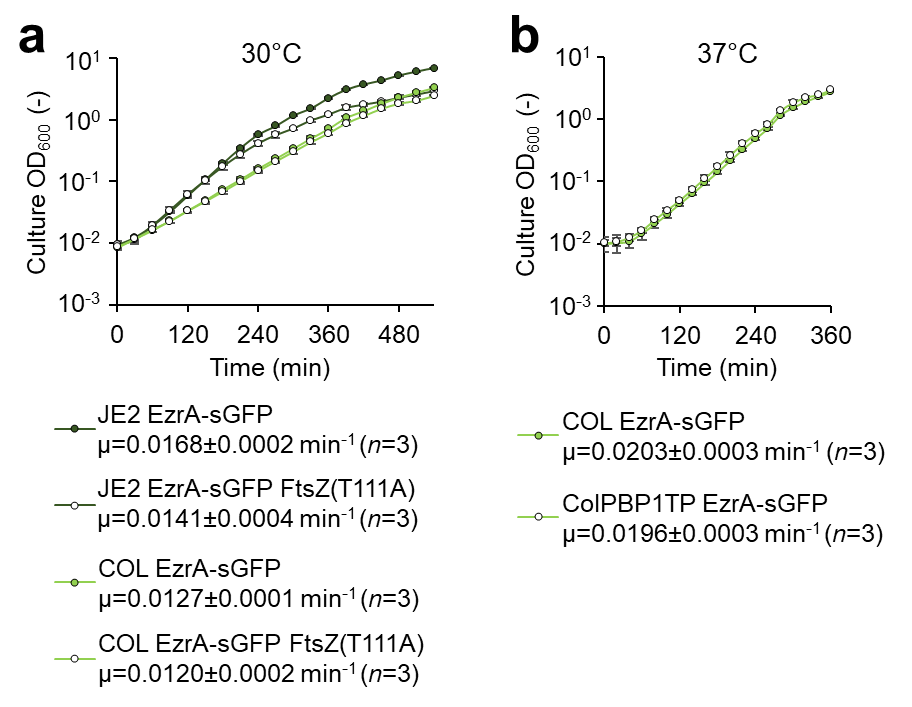

**Figure S3. Growth curves of *S. aureus* FtsZ GTPase and PBP1 TPase mutant strains.** Culture growth curves of indicated strains in TSB rich medium at indicated temperatures. The OD_600_ was recorded every 30 min (**a**) or 20 min (**b**). Error bars represent the standard deviations for the mean from three biological replicates. Growth rate (µ) was calculated for cells in mid-exponential phase.

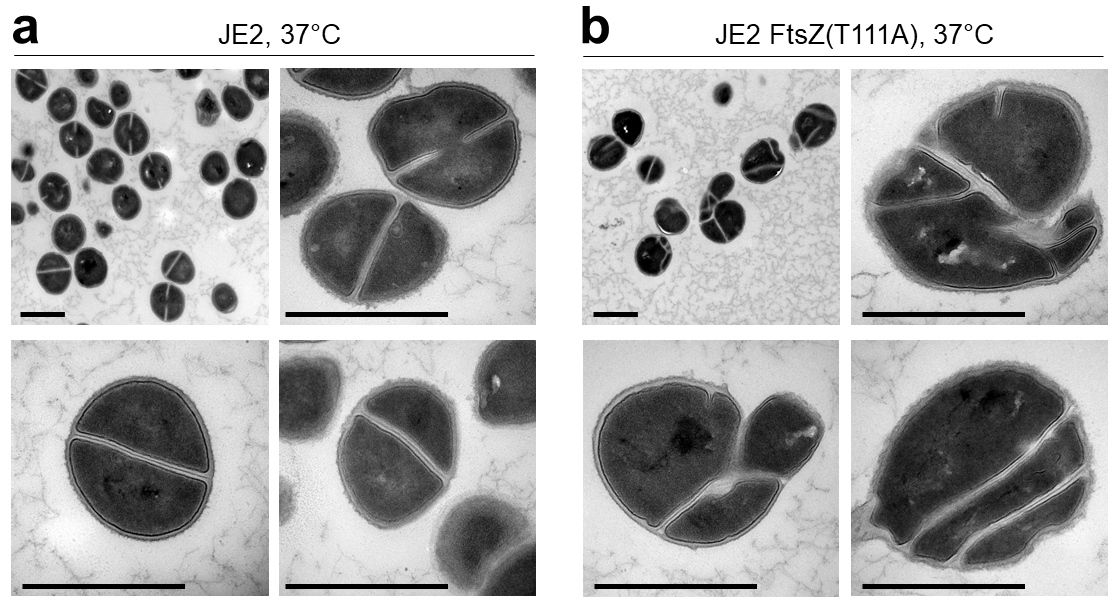

**Figure S4. The FtsZ GTPase mutation T111A causes cell morphology defects in *S. aureus*.** Representative electron micrographs of JE2 wild-type and mutant derivative FtsZ(T111A) strains grown to mid-exponential phase in TSB rich medium at 37°C. Scale bars, 1 µm.

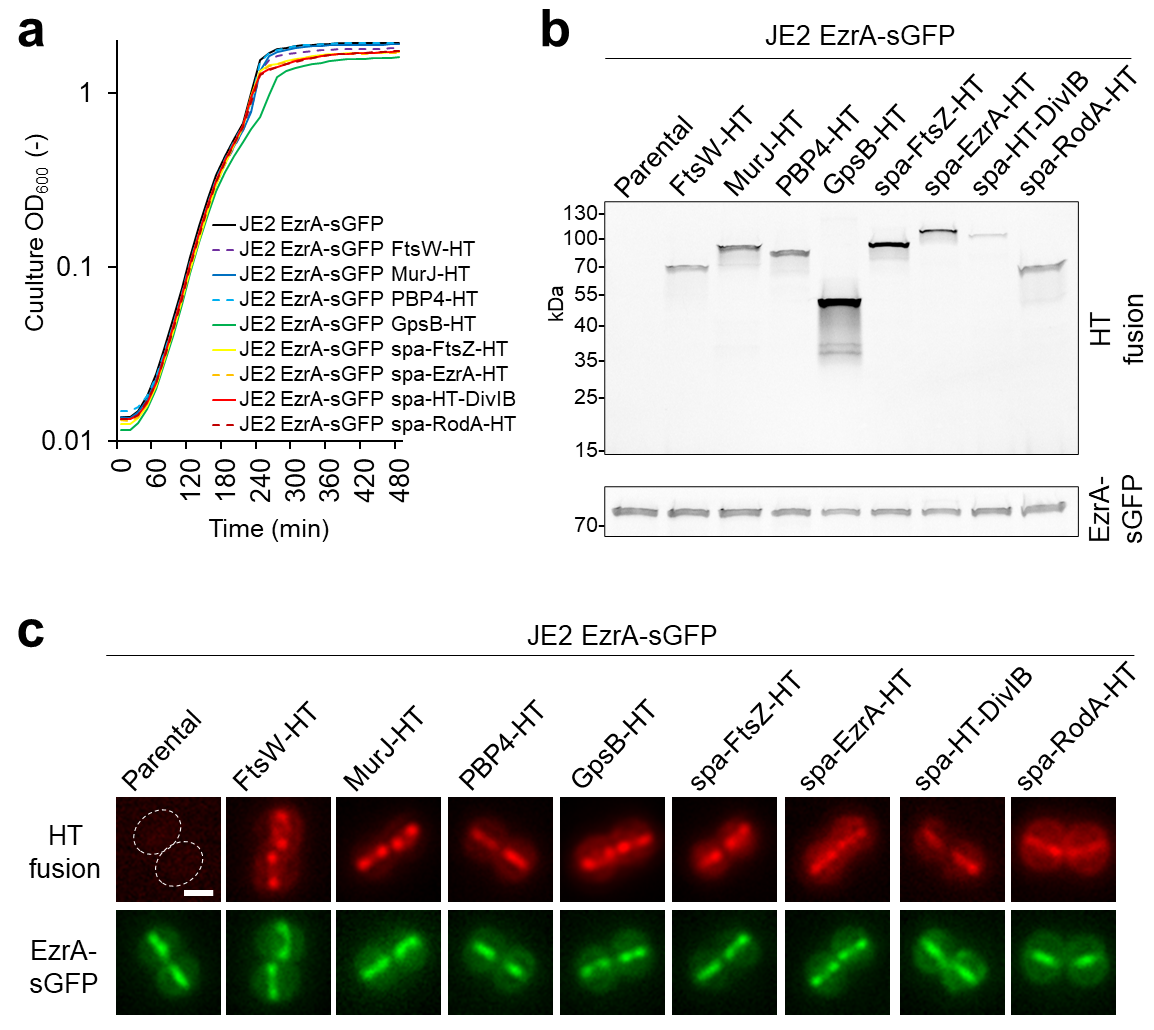

**Figure S5. Halo-tag fusions to cell division and cell wall-related proteins produced in *S. aureus* are non-toxic and enriched at mid-cell. a-d,** Strains indicated in panel **a** were grown shaking in TSB rich medium at 37°C. Cells producing EzrA-HT or HT-DivIB were grown in presence of 0.5 ng/ml Atc or 0.5 mM IPTG, respectively. **a,** Growth curves recorded in 96-well plate format and obtained from six biological replicates. **b,c,** Fluorescent protein gel (**b**) and representative epifluorescence micrographs (**c**) of cells grown to mid-exponential phase and labelled with 500 nM JF549-HTL. Theoretical molecular weights (in kDa): FtsW-HT, 79.6; MurJ-HT, 96.7; PBP4-HT, 83.0; GpsB-HT, 47.2; FtsZ-HT, 75.7; EzrA-HT, 101.0; HT-DivIB, 84.5; RodA-HT, 78.7; EzrA-sGFP, 93.6. Dashed lines in the negative control indicate cell outlines inferred from the corresponding phase contrast image. Scale bar, 0.5 µm.

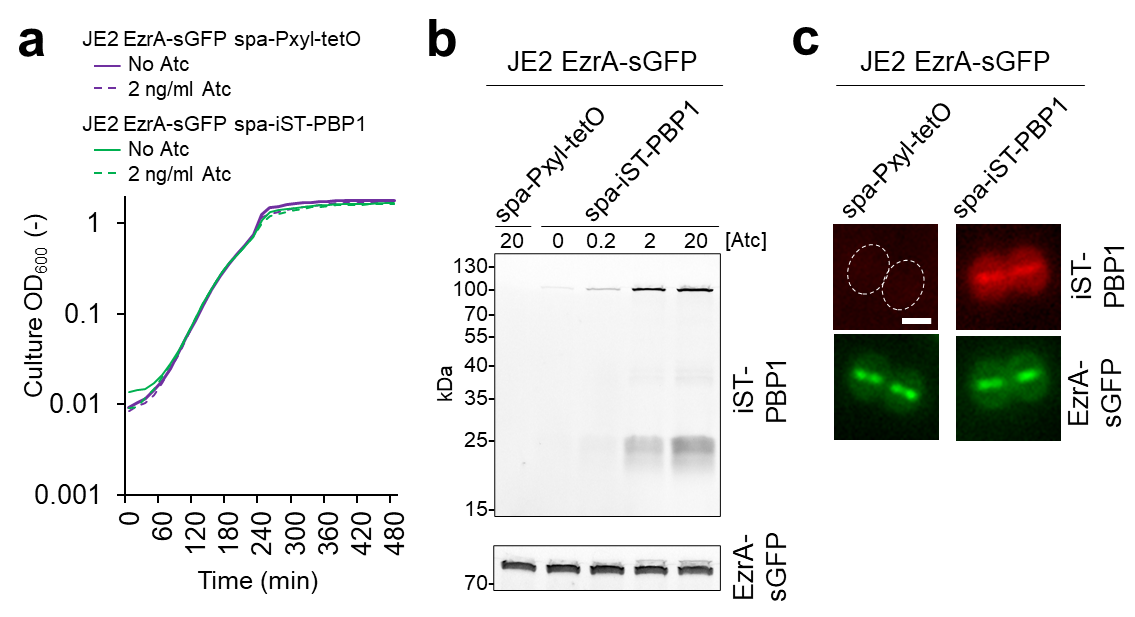

**Figure S6. A Snap-tag fusion to PBP1 produced in *S. aureus* is non-toxic and enriched at mid-cell. a-d,** Strains indicated in panel **a** were grown shaking in TSB rich medium at 37°C and where indicated in presence of Anhydrotetracycline (Atc) to induce gene expression from the heterologous xylose promoter (P*xyl-tetO*). **a,** Growth curves recorded in 96-well plate format and obtained from six biological replicates. **b,c,** Fluorescent protein gel (**b**) and epifluorescence micrographs (**c**) of cells grown to mid-exponential phase and labelled with 500 nM JF549-cpSTL. The theoretical molecular weights of iST-PBP1 and EzrA-sGFP are 103.7 and 93.6 kDa, respectively. Atc concentration is given in ng/ml. Dashed lines in the negative control indicate cell outlines inferred from the corresponding phase contrast image. Scale bar, 0.5 µm.

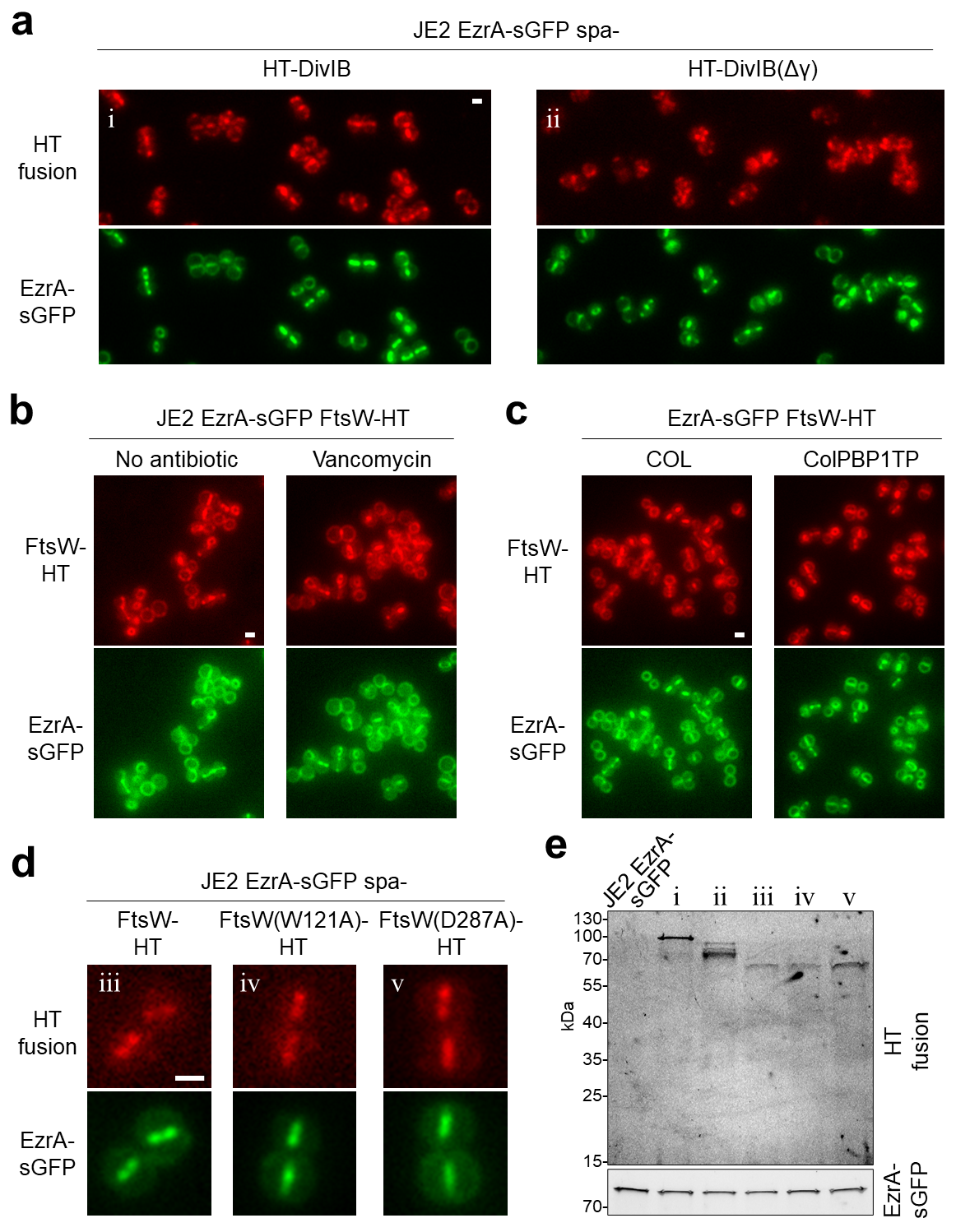

**Figure S7. Localization of DivIB and FtsW variants in wild-type cells or in cells with perturbed peptidoglycan synthesis. a,** Representative epifluorescence micrographs of cells producing HT fused to DivIB wild-type or DivIB lacking its C-terminal γ domain from the ectopic *spa* locus. Indicated strains were grown in TSB rich medium supplemented with 0.5 mM IPTG at 37°C. Cells were labelled with 500 nM JF549-HTL after reaching mid-exponential growth phase. **b,c,** Representative epifluorescence micrographs of cells producing FtsW-HT from its native genomic locus in the indicated genetic backgrounds. Exponentially growing cells were incubated with 500 nM JF549-HTL and with or without 2 µg/ml vancomycin for 20 min at 37°C. Scale bars, 0.5 µm. **d,** Representative epifluorescence micrographs of cells producing FtsW-HT wild-type or active-site mutant derivatives W121A and D287A from the ectopic *spa* locus. Indicated strains were grown in TSB rich medium supplemented with 0.2 ng/ml Atc at 37°C. Cells were labelled with 500 nM JF549-HTL after reaching mid-exponential growth phase. **e,** Fluorescent protein gel for strains indicated by numbers in panels **a** and **d**. Theoretical molecular weights (in kDa): HT-DivIB, 84.5; HT-DivIB(Δγ), 77.1; FtsW-HT, 79.6; EzrA-sGFP, 93.6.

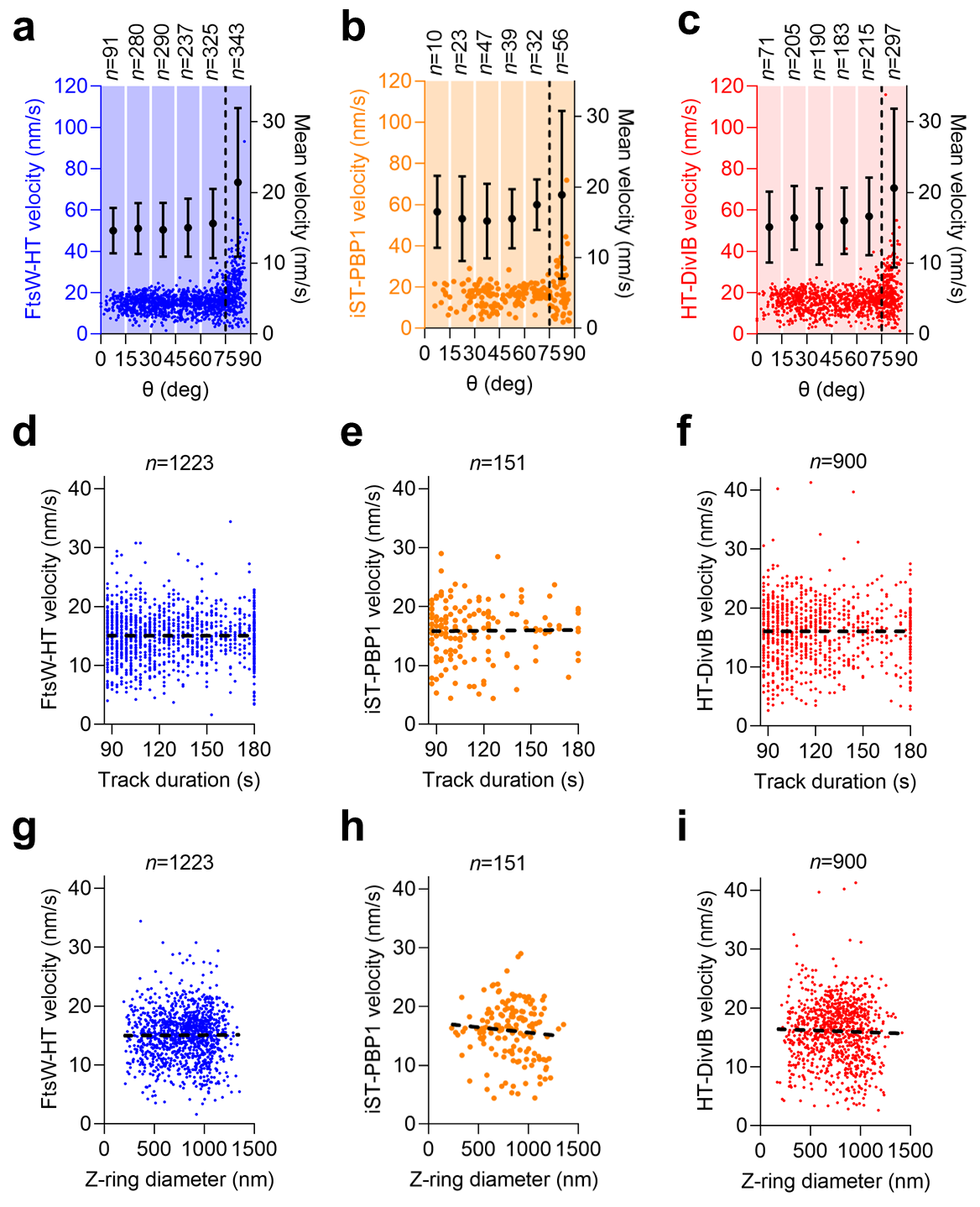

**Figure S8. FtsW, PBP1 and DivIB velocities do not correlate with track duration or cell division stage.** FtsW-HT (**a,d,g**), iST-PBP1 (**b,e,h**) and HT-DivIB (**c,f,i**) single-molecule velocity as a function of the angle between the imaging and cell division planes (Θ) (**a-c**), track duration (**d-f**) and EzrA-sGFP ring diameter (**g-i**). Each point in a graph corresponds to the calculated average velocity of a trajectory. Cells were grown in TSB rich medium at 37°C and in presence of 0.5 mM IPTG (HT-DivIB) or 2 ng/ml Atc (iST-PBP1). The cut-off for Θ at 75° (**a-c**) and simple linear regressions (**d-i**) are indicated as dashed lines. Black points and error bars represent the means and standard deviations for average track velocities in the indicated range of Θ.

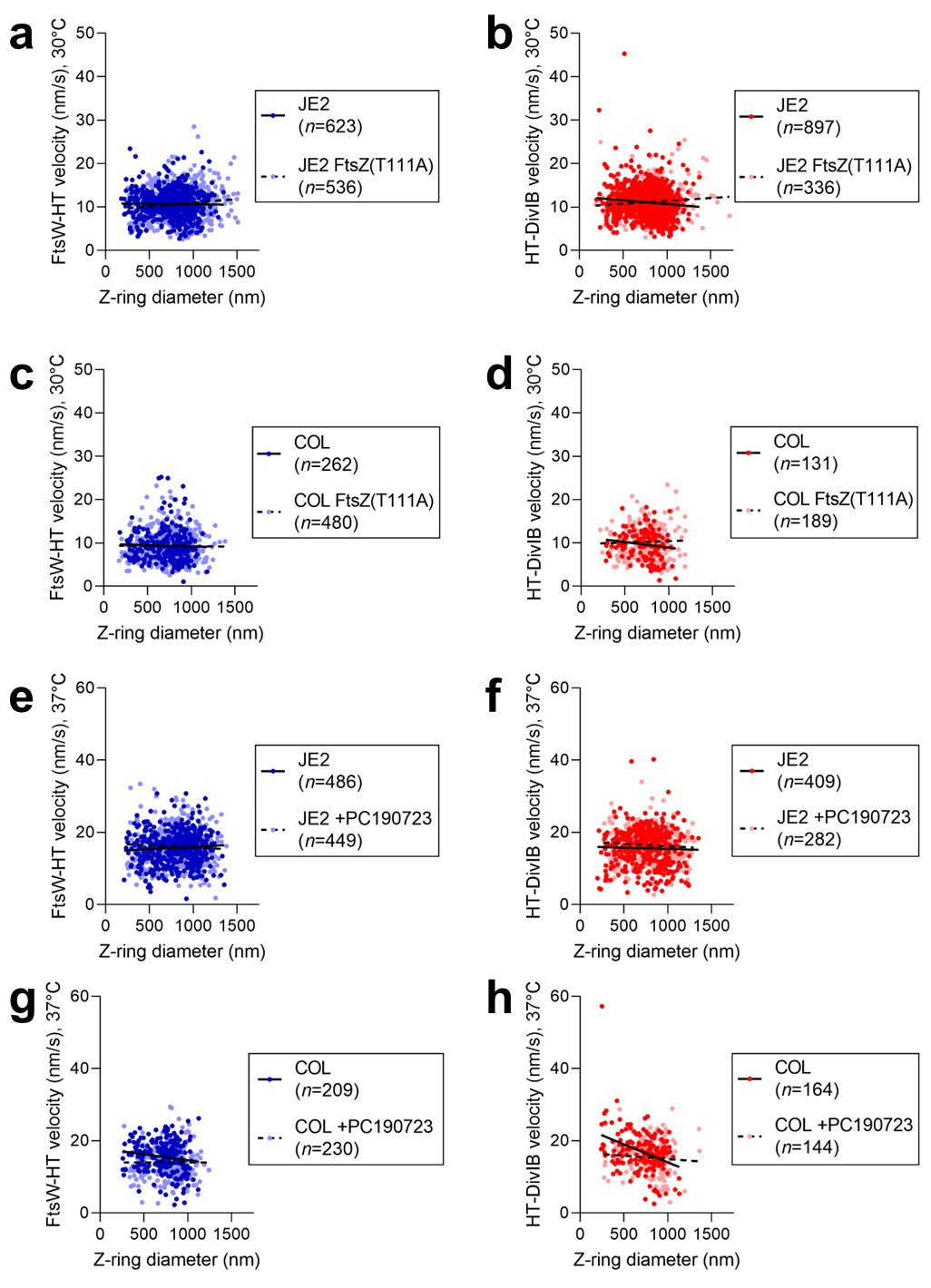

**Figure S9. FtsW and DivIB velocities remain unchanged when FtsZ treadmilling is impaired at any stage of cell division.** FtsW-HT (**a,c,e,g**) and HT-DivIB (**b,d,f,h**) single-molecule velocity as a function of EzrA-sGFP ring diameter in cells containing FtsZ mutation T111A or treated with 5 µg/ml PC190723 for 2 min. Each point in a graph corresponds to the calculated average velocity of a trajectory. Cells were grown in TSB rich medium at 30°C (**a-d**) or at 37°C (**e-h**), and in presence of 0.5 mM IPTG (**b,d,f,h**). Simple linear regressions are indicated as continuous and dashed lines.

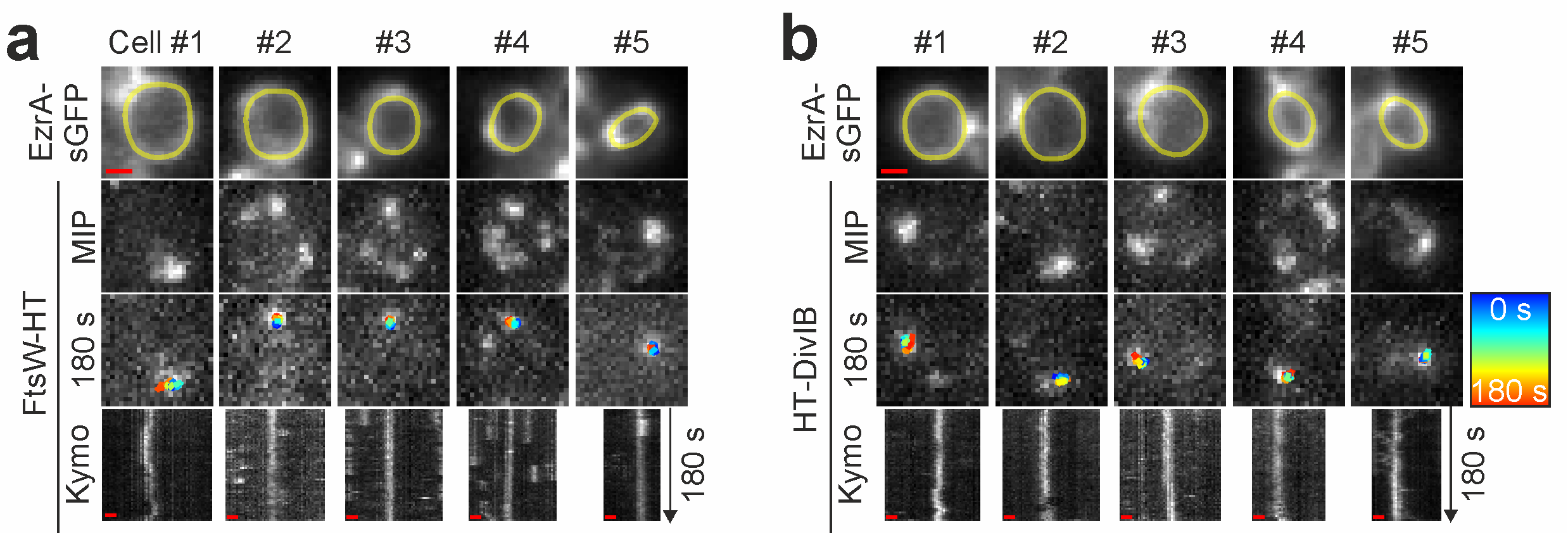

**Figure S10. Directional movement of FtsW and DivIB is stopped by vancomycin. a,b,** Representative epifluorescence micrographs of JE2 EzrA-sGFP producing FtsW-HT (**a**) or HT-DivIB (**b**). Cells producing HT-DivIB were grown in presence of 0.5 mM IPTG to induce gene expression from the ectopic *spa* locus. Cells were treated with 2 µg/ml vancomycin for 20 min at 37°C and sparsely labelled with the fluorescent ligand JF549-HTL to visualize single molecules of FtsW-HT and HT-DivIB. Five independent cells are shown in each panel. Single-molecule images acquired in the last frame of a 180-s time series are overlayed with tracks, where blue (0 s) to red (180 s) indicates trajectory time. Space-time kymographs were generated by extracting fluorescence intensity values from FtsW-HT and HT-DivIB images along yellow lines drawn over corresponding EzrA-sGFP rings acquired in the last frame of each time series. MIP, maximum intensity projection. Scale bars, 0.5 µm.

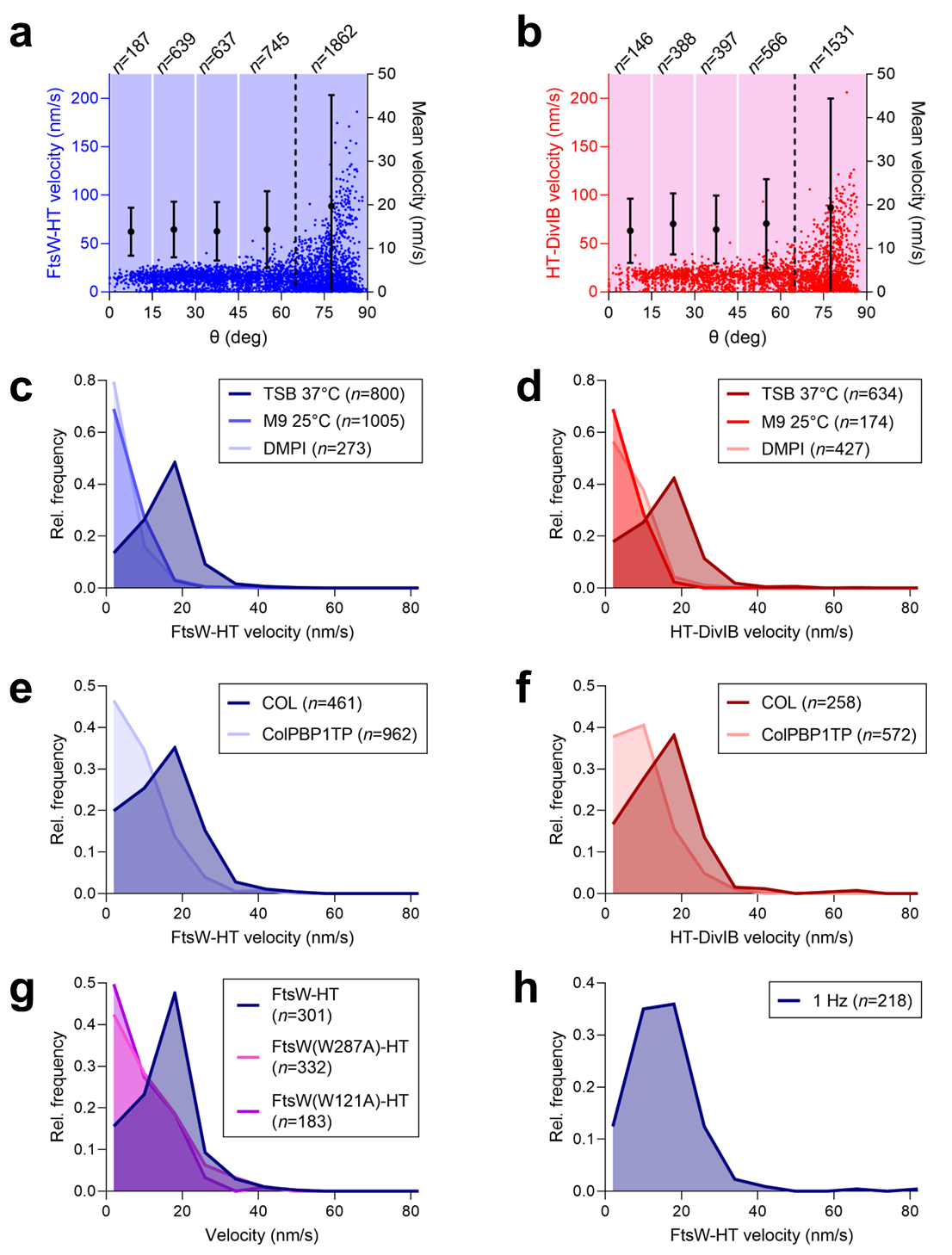

**Figure S11. FtsW and DivIB sectional velocities show a unimodal distribution. a,b,** FtsW-HT (**a**) and HT-DivIB (**b**) single-molecule velocity as a function of the angle between the imaging and cell division planes (Θ). Each point in a graph corresponds to the calculated velocity in each section of a trajectory. The cut-off for Θ at 65° is indicated as dashed line. Black points and error bars represent the means and standard deviations for velocities in each section in the indicated range of Θ. Data was obtained for JE2 EzrA-sGFP grown in TSB rich medium at 37°C. **c-h,** Histograms depicting the velocity distribution for FtsW-HT (**c,e,g,h**) and HT-DivIB (**d,f**) determined in cells of strain JE2 EzrA-sGFP grown in the indicated conditions. Each data point in the histogram corresponds to the velocity of a single section. Bin width, 8. Center of first/last bin, 2/82. Unless otherwise specified, the frame rate for image acquisition was 0.33 Hz.

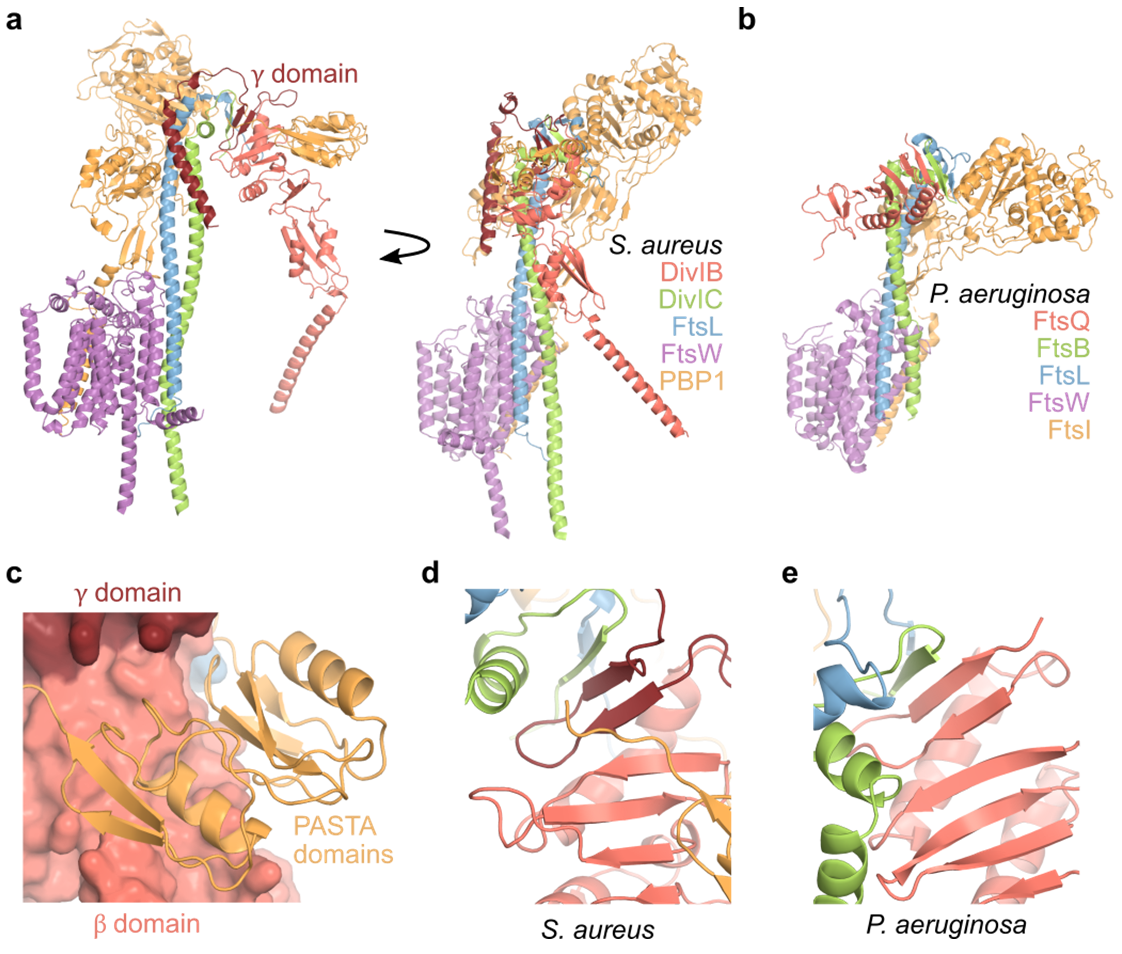

**Figure S12. Structure prediction of a putative pentameric complex composed of the *S. aureus* proteins FtsW, PBP1, DivIB, DivIC and FtsL.** **a,** Alphafold-multimer model of the complex formed by FtsW (purple), PBP1 (orange), DivIB (red), DivIC (green) and FtsL (blue) from *S. aureus*. DivIB γ-domain residues 373-439 are shown in a darker red color. Terminal residues with low local prediction confidence are not shown (pLDDT below 50) with the exception of DivIB C-terminal residues and PBP1 residues 590–594 between the TPase and PASTA domains. **b,** Comparison to the cryo-EM structure of the orthologous *P. aeruginosa* complex (PDB 8BH1) after alignment to FtsW shows a similar predicted arrangement of domains and a similar tilt of the TPase domain to that observed between predicted and experimental structures of the divisome complex ^11,12^. **c,** C-terminal PASTA domains in PBP1 are predicted to interact primarily with the DivIB β domain. **d,e,** Comparison of the local structures after alignment to γ-domain residues shows that the *S. aureus* γ domain is predicted to contribute two strands to a β sheet linking DivIB, DivIC, and FtsL, as was observed experimentally for *P. aeruginosa*. Relative orientations of DivIC and FtsL helices differ in the *S. aureus* prediction.

**Supplementary Movie 1.** Treadmilling in FtsZ(T111A) mutant and parental strains grown at 30°C.

**Supplementary Movie 2.** Septum constriction in FtsZ(T111A) mutant and parental strains grown at 30°C.

**Supplementary Movie 3.** Treadmilling in COL derivative strains in the absence and presence of indicated antibiotics and grown at 37°C.

**Supplementary Movie 4.** Septum constriction in COL derivative strains in the absence and presence of indicated antibiotics and grown at 37°C.

**Supplementary Movie 5.** Single-molecule tracking of FtsW-HT and HT-DivIB in JE2 derivative strains in the absence and presence of indicated antibiotics and grown at 37°C.

**Supplementary Movie 6.** Animation illustrating the single-molecule tracking data analysis for molecules moving directionally around the division site of coccoid cells placed on a microscope slide.

**Supplementary Movie 7.** Single-molecule tracking of FtsW-HT and HT-DivIB in FtsZ(T111A) mutant and parental strains grown at 30°C.

**Supplementary Movie 8.** Single-molecule tracking of FtsW-HT and HT-DivIB in COL and ColPBP1TP strains grown at 37°C.
